# A novel *Smg6* mouse model reveals regulation of circadian period and daily CRY2 accumulation through the nonsense-mediated mRNA decay pathway

**DOI:** 10.1101/2022.07.01.498406

**Authors:** Georgia Katsioudi, René Dreos, Enes S. Arpa, Sevasti Gaspari, Angelica Liechti, Miho Sato, Christian H. Gabriel, Achim Kramer, Steven A. Brown, David Gatfield

## Abstract

Nonsense-mediated mRNA decay (NMD) has been intensively studied as a surveillance pathway that degrades erroneous transcripts arising from mutations or RNA processing errors. While additional roles in controlling regular mRNA stability have emerged, possible functions in mammalian physiology *in vivo* have remained unclear. Here, we report a novel conditional mouse allele that allows converting the NMD effector nuclease SMG6 from wild-type to nuclease domain-mutant protein. We analyzed how NMD downregulation affects the function of the circadian clock, a system known to require rapid mRNA turnover. We uncover strong lengthening of free-running circadian periods for liver and fibroblast clocks, and direct NMD regulation of *Cry2* mRNA, encoding a key transcriptional repressor within the rhythm-generating feedback loop. In the entrained livers of *Smg6* mutant animals we reveal transcriptome-wide alterations in daily mRNA accumulation patterns, altogether expanding the known scope of NMD regulation in mammalian gene expression and physiology.

## Introduction

Nonsense-mediated mRNA decay (NMD) functions as an important surveillance pathway to reduce gene expression errors that arise from mutations or mis-splicing and that are recognized due to aberrant translation termination on “premature translation termination codons” (PTCs) (reviewed in ^1,2^). In mammals, PTCs are defined due to their position relative to an exon-junction complex (EJC), a multiprotein assembly that is deposited on mRNAs during splicing and removed from the transcript by the passage of translating ribosomes. Termination upstream of an EJC identifies the stop codon as aberrant, promoting the formation of an NMD factor complex comprising several UPF (up-frameshift) and SMG (suppressor with morphogenetic effects on genitalia) proteins. Briefly, interactions between UPF1, UPF2 and UPF3 proteins trigger UPF1 phosphorylation by the kinase SMG1. Phosphorylated UPF1 further recruits SMG5, SMG6 and SMG7, which are involved in executing the actual mRNA degradation step. Previous models suggested two distinct, redundant branches for decay involving SMG5-SMG7 (that can recruit general, non-NMD-specific exonucleases) or SMG6 (an NMD-specific endonuclease); recent evidence, however, argues for mechanistic overlap ^3^, and a linear pathway involving decay “licensing” through SMG5-SMG7 followed by SMG6-mediated endonucleolytic cleavage has been proposed as the main mechanism of how mRNA decay is carried out ^4^.

Early transcriptome-wide analyses already noted that in addition to NMD activity on aberrant transcripts, the pathway can be co-opted for the decay of regular, physiological mRNAs as well ^5^. Most of the initially identified NMD-activating features on endogenous transcripts are in line with the above rules for PTC definition - e.g.: introns in 3’ untranslated regions (UTRs); translated upstream open reading frames (uORFs) in 5’ UTRs; selenocysteine codons that are interpreted as stop codons - yet later studies showed that long 3’ UTRs can activate NMD *per se*, in the absence of a downstream splice junction ^6,7^. The generality of a “3’ UTR length rule” has, however, been questioned recently in a nanopore sequencing-based study that (after removing the transcripts from the analysis for which there was evidence for splicing in the 3’ UTR) found no predictive value of 3’ UTR length for NMD regulation ^8^. Independently of which mechanisms trigger NMD on non-classical NMD substrates, it has been proposed that the expression of up to 20-40% of genes is directly or indirectly affected when NMD is inactivated in mammalian cell lines ^4,8^, and it is tempting to speculate that the pathway may have extensive functions in regular gene expression control ^2^. It is largely unknown whether this regulatory potential of NMD extends to the intact organ and living organism *in vivo*, and if so, which specific molecular and physiological pathways it controls.

Certain physiological processes are particularly reliant on rapid, well-controlled RNA turnover; conceivably, co-opting NMD could thus be especially opportune. In this respect, the circadian clock stands out as an important functional system that controls daily rhythms in transcription, mRNA and protein abundances, affecting thousands of genes across the organism and controlling daily changes in behavior, physiology and metabolism (reviewed in ^9^). In the mammalian body, the circadian system is hierarchically organized with a master clock in the brain’s suprachiasmatic nucleus (SCN) that synchronizes peripheral clocks that operate in most cell types and that are responsible for driving cellular rhythmic gene expression programs. Across cell types, clocks have a similar molecular architecture, with a core clock mechanism that generates gene expression oscillations through transcription factors that interact in negative feedback loops. In the main loop, BMAL1:CLOCK (BMAL1:NPAS2 in neurons) function as the main activators and bind to E-box enhancers in their target genes, which include the *Period* (*Per1, 2, 3*) and *Cryptochrome* (*Cry1, 2*) genes. Negative feedback is achieved when PER:CRY complexes translocate to the nucleus and repress their own transcription by inhibiting BMAL1:CLOCK. PER and CRY protein degradation temporally limits the repressive activity, eventually allowing a new cycle to ensue. Conceivably, the rapid decay of *Per* and *Cry* mRNAs is critical for this mechanism - as a means of restricting PER:CRY biosynthesis and availability in time - yet the responsible decay pathways remain poorly investigated. Additional feedback mechanisms (in particular involving nuclear receptors of the REV-ERB/ROR families) interlock with the above main feedback loop and confer both robustness and plasticity to this system (reviewed in ^10^). Through the numerous rhythmic transcriptional activities that are generated through this clockwork, rhythmic mRNA production is driven at hundreds to thousands of downstream, clock-controlled genes (CCGs). The stability of CCG transcripts critically determines to what extent their initial transcriptional rhythm is propagated to the mRNA and protein abundance levels. Mechanisms that have been implicated in post-transcriptionally regulating rhythmic mRNAs in mammals include miRNA-mediated mRNA decay ^11^ and regulated deadenylation ^12^. With regard to a possible involvement of NMD, first evidence for roles in the circadian system has been reported from fungi, plants and flies ^13-15^, but how NMD globally shapes rhythmic transcriptomes, let alone in a mammalian organism *in vivo*, is still unknown.

In this study, we have comprehensively investigated the role of NMD in the mammalian circadian system *in vivo*. Using a novel conditional NMD loss-of-function mouse model, we uncover that NMD is directly implicated in regulating the circadian period of peripheral clocks. We identify *Cry2* as a direct NMD target and further determine how the hepatic diurnal transcriptome is rewired in the absence of a functional NMD pathway. Our new mouse model and findings on circadian regulation provide important conceptual advances on *in vivo* functions of NMD and reveal a novel mechanism of post-transcriptional gene expression regulation that acts in the mammalian core clock.

## Results

### A novel conditional NMD loss-of-function allele based on SMG6 mutated in its nuclease domain

To inactivate NMD *in vivo* we generated mice in which we could conditionally recombine *Smg6*^*flox*^ to *Smg6*^*mut*^ (**Fig. 1A**), i.e. from an allele encoding wild-type SMG6 protein to a version specifically point-mutated at two of the three highly conserved aspartic acid (D) residues of the catalytic triade of the protein’s PIN (PilT N-terminus) nuclease domain ^16^ (**Fig. 1B**). We chose this strategy over a full gene knockout because NMD factors, including SMG6, carry additional functions in telomere and genome stability ^17^. These functions have been shown to be selectively maintained by expressing an NMD-inactive SMG6 protein lacking its nuclease domain ^18^. We first validated our genetic model in primary tail fibroblasts from homozygous *Smg6*^*flox/flox*^ and *Smg6*^*+/+*^ littermate mice that we stably transduced with a retroviral vector expressing tamoxifen-activatable CreERT2 (**Fig. 1C**). *Smg6*^*flox/flox*^ cells specifically and efficiently recombined to *Smg6*^*mut/mut*^ by addition of 4-hydroxytamoxifen (4-OHT) to the culture medium (**Fig. 1D**). In these cells, a lentiviral luciferase reporter carrying an intron in its 3’ UTR became upregulated, as expected for an inactive NMD pathway (**Fig. 1E**). We next used RNA-seq on 4-OHT-treated and -untreated cells of both genotypes to analyze gene expression changes transcriptome-wide. Our method, based on random priming of rRNA-depleted total RNA, allowed for the quantification of both mRNA (exon-mapping reads) and pre-mRNA abundances (intron-mapping reads), the latter serving as a proxy for gene transcription rates ^11,19,20^. In analogy to previous studies ^11,19^ we used mRNA/pre-mRNA ratios to estimate mRNA stability changes between NMD-inactive and control cells, and to distinguish them from secondary effects involving altered transcription rates. Our analyses revealed a shift to higher mRNA/pre-mRNA ratios (more stable mRNAs) specifically in NMD-inactive (*Smg6*^*flox/flox*^ + 4-OHT) cells (**Fig. 1F**). Two transcript groups that were particularly affected, as predicted, were genes with known, annotated NMD-sensitive mRNA isoforms (according to Ensembl annotations) and with retained introns (**Fig. 1G**). Visual inspection of individual examples further validated these findings, as shown for *Hnrnpl* and *Srsf11*, for which a specific up-regulation of NMD isoform-specific exons in the mutants was evident (**Fig. 1H-I**). Transcriptome-wide differential expression analysis at the exon level indicated that apart from NMD-annotated isoforms and retained introns, hundreds of constitutive exons from canonical mRNAs (i.e., without annotated NMD isoforms) increased in abundance under *Smg6* mutant conditions, pointing to widespread NMD regulation of the transcriptome (**Fig. 1J**). We next analyzed if specific transcript features correlated with *Smg6* mutation-dependent changes in mRNA/pre-mRNA ratios. As expected for potential NMD substrates, the transcripts that were most strongly affected were low expressed in control cells (**Fig. 1K**). 5’ UTR length (which correlates with uORF content ^21^) was weakly, though significantly, associated with increased mRNA/pre-mRNA ratios (**Fig. 1L**), suggesting that translated uORFs may contribute as an NMD-activating feature to endogenous mRNA upregulation in *Smg6* mutants. Stronger correlations were observed with the lengths of the CDS (**Fig. 1M**) and 3’ UTRs (**Fig. 1N**). The latter association is consistent with the ability of long 3’ UTRs to function as NMD-activating features. Altogether these associations match those observed for other NMD loss-of-function models, e.g. in Hela cells subjected to *Upf1* knockdown ^22^. Taken together, we concluded that our genetic model based on mutant *Smg6* was suitable to analyze endogenous targets and functions of the NMD pathway.

**Figure 1.**
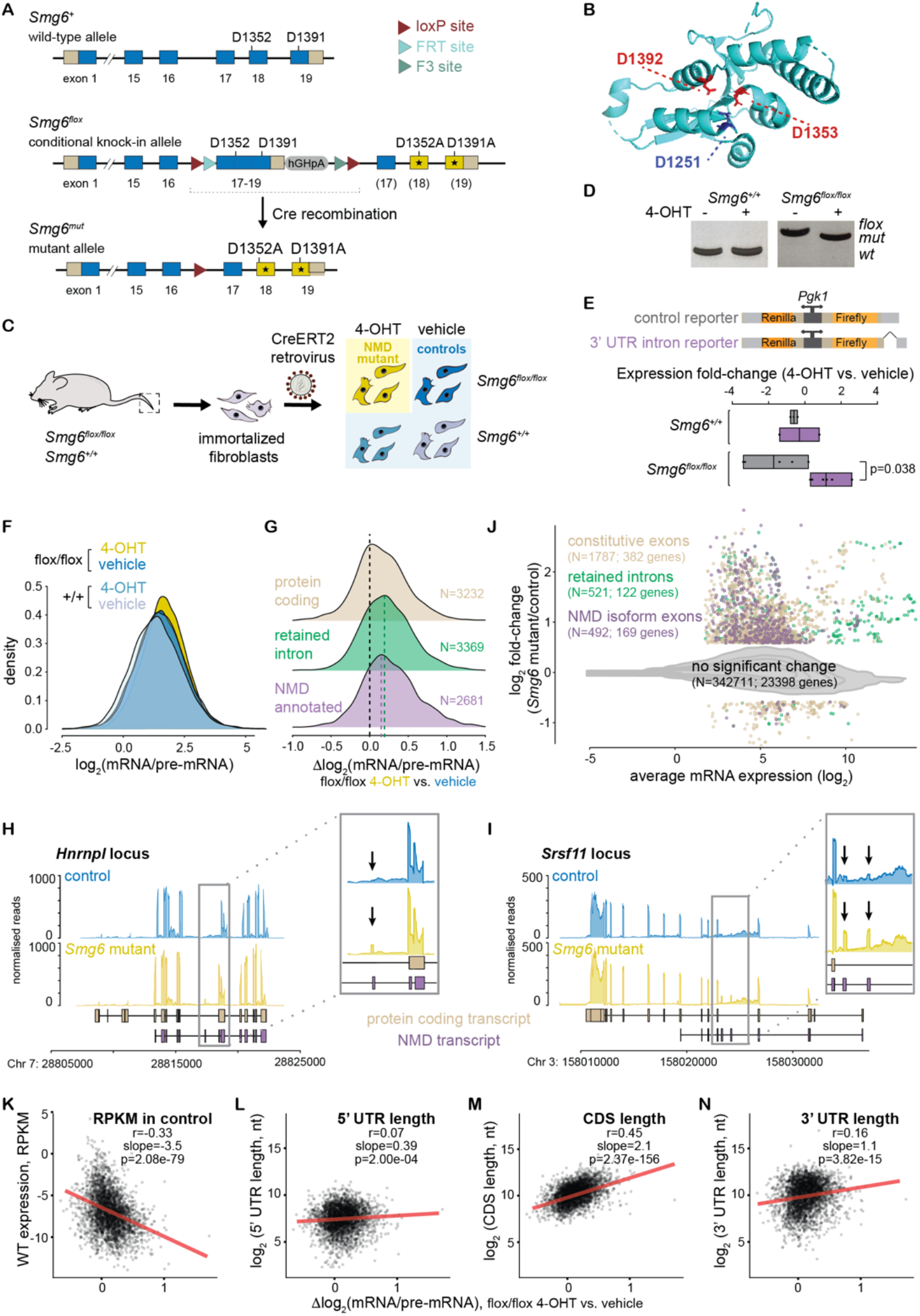
A novel conditional NMD loss-of-function allele based on PIN nuclease domain mutant *Smg6*. **A**. Schematic of the genetic model. *Smg6*^*flox*^ expresses wild-type SMG6 protein encoded by the blue exons; after Cre-mediated recombination to *Smg6*^*mut*^, point-mutated exons 18 and 19 (yellow) lead to expression of mutant SMG6 (D1352A, D1391A). **B**. The mutated aspartic acid residues are within the catalytic triade of the PIN nuclease domain, shown in the structure of the human protein (PDB accession 2HWW; ^16^). **C**. For cellular studies, tail fibroblasts from adult male mice (*Smg6*^*flox*^ and wild-type littermates) were cultured until spontaneous immortalization, and tamoxifen-activatable CreERT2 expression was achieved by a retrovirus. Upon 4-hydroxytamoxifen (4-OHT) treatment, NMD mutants (yellow) were compared to different control cells (shades of blue). **D**. PCR-based genotyping of genomic DNA extracted from cells depicted in C. indicates efficient recombination upon 4-OHT treatment. **E**. A luciferase reporter containing an intron in the 3’ UTR is upregulated in 4-OHT-treated *Smg6*^*flox/flox*^ cells, as expected under NMD-inactive conditions. N=2-6 plates/group, adjusted p=0.0038; multiple Student’s t-test. **F**. Density plot showing transcriptome-wide mRNA/pre-mRNA ratio distributions calculated from RNA-seq, in NMD-inactive (yellow) vs. control cells. **G**. The difference in mRNA/pre-mRNA ratios between NMD-inactivated (*Smg6*^*flox/flox*^ + 4-OHT) and control cells (*Smg6*^*flox/flox*^ + vehicle) is consistent with higher stability of annotated NMD substrates (purple, N=2681) and transcripts with retained introns (green, N=3369). Moreover, the broad distribution and shift to positive values for non-NMD-annotated protein coding transcripts (beige, N=3232) is indicative of transcriptome-wide NMD regulation. **H**. Read coverage on the *Hnrnpl1* and **I**. *Srsf11* loci indicates the specific upregulation of transcript isoforms that are NMD-annotated (purple) and that can be identified via specific exons (marked by arrows in insets). **J**. Differential expression analysis at the exon level, comparing *Smg6*^*flox/flox*^ + 4-OHT vs. *Smg6*^*flox/flox*^ + vehicle conditions, reveals significant upregulation of NMD-annotated exons (purple; N=492; 169 genes), retained introns (green; N=521; 122 genes), and a sizeable number of constitutive exons (beige; N=1787, 382 genes), suggestive of NMD regulating many protein coding genes. **K**. Correlation analysis between mRNA/pre-mRNA ratio change upon NMD activation and expression levels in wild-type cells shows significant anticorrelation. The lengths of **L**. the 5’ UTR, **M**. the CDS and **N**. the 3’ UTR are all positively correlated with mRNA/pre-mRNA ratio change upon NMD inactivation. Pearson correlation coefficient (r), slope and p-values were calculated by a linear model.

### NMD inactivation lengthens free-running circadian periods in fibroblasts and in liver

We next investigated how mutant *Smg6* affected the circadian clock. First, we stably transfected the above fibroblasts with a circadian reporter gene, *Dbp-Luciferase* ^23^, and recorded their free-running circadian rhythms upon NMD inactivation with 4-OHT. Briefly, we synchronized the cellular oscillators using temperature cycles ^24^, released them at 37°C, and continued real-time bioluminescence recordings for an additional 5 days under constant conditions (**Fig. 2A**). These experiments revealed a lengthening of the free-running circadian period in NMD-deficient cells by ca. 1.5 hours (**Fig. 2B**). We next wished to corroborate a potential period phenotype using an alternative peripheral clock model that was more relevant for circadian physiology and functions *in vivo*. We thus crossed into the *Smg6*^*flox*^ mouse line a hepatocyte-specific *CreERT2* (driven from the *Albumin* locus ^25^) and a circadian reporter, *mPer2::Luc* ^26^. After intraperitoneal tamoxifen injections into young adult mice, animals were sacrificed 4 weeks later, a time at which highly efficient recombination to *Smg6*^*mut*^ had taken place (**Fig. 2E**). We then prepared organotypic slices (tissue explants) for real-time recording of luciferase rhythms *ex vivo* (**Fig. 2C**). In these experiments we observed a strong and specific period lengthening by almost 3 hours in liver explants from animals with inactivated NMD (tamoxifen-treated *Smg6*^*flox/flox*^ mice) as compared to livers from identically treated littermate animals of the control genotype (**Fig. 2D**). As an additional specificity control, we recorded kidney explant rhythms from the same animals. Free-running periods were generally longer in this organ, as reported previously ^26^, yet we did not observe any differences between genotypes (**Fig. 2D**), in line with the hepatocyte-specificity of CreERT2 expression in our genetic model.

**Figure 2.**
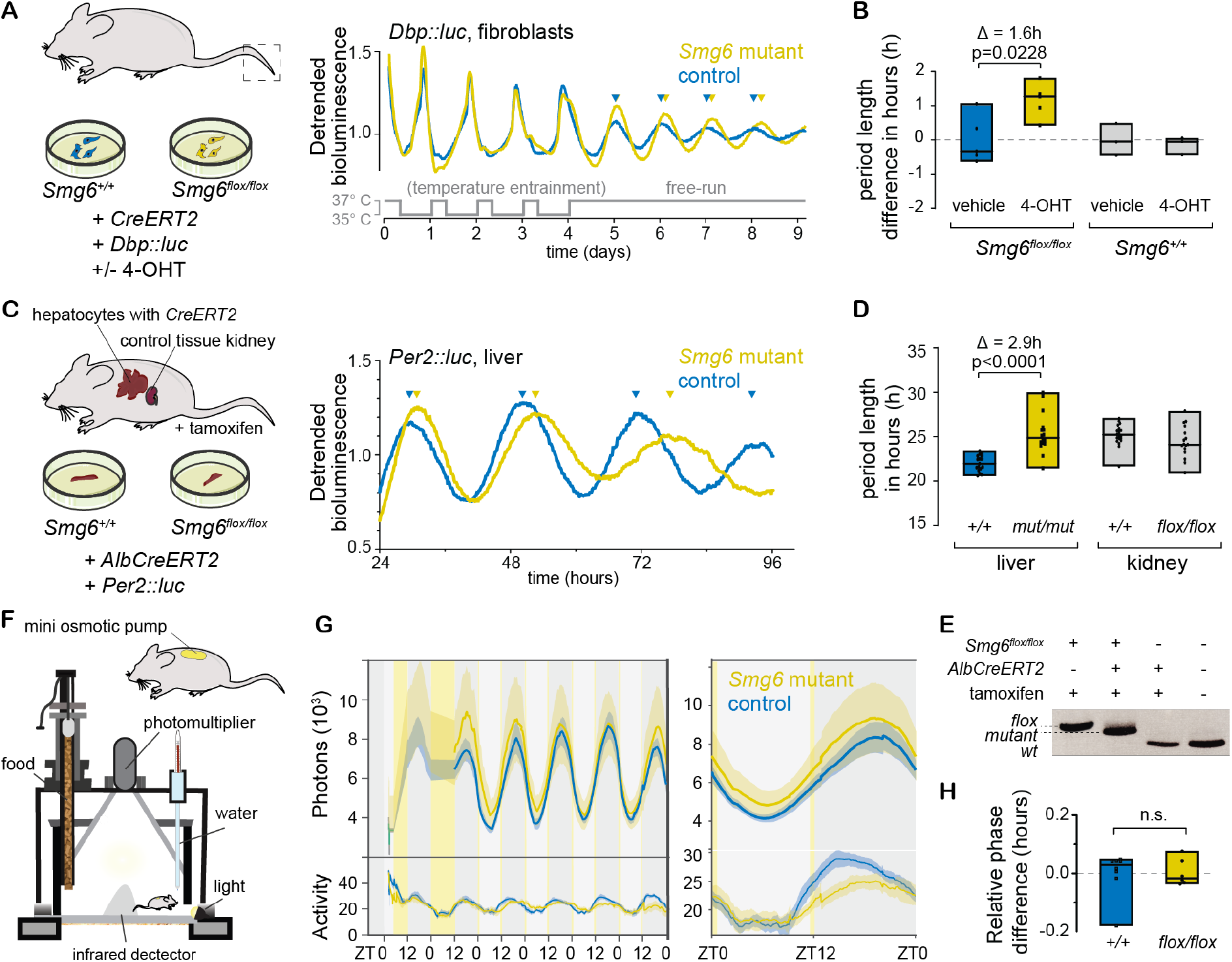
NMD inactivation through *Smg6* mutation lengthens free-running circadian periods. **A**. Real-time recording of bioluminescence rhythms in immortalized fibroblasts of genotypes *Smg6*^*flox/flox*^ and *Smg6*^*+/+*^ (both transduced with CreERT2 retrovirus and circadian reporter *Dbp::Luc*); after temperature-entrainment (24-h periodic square wave; 35°C-37°C), cells were released into free-running conditions (37°C) from day 4. Representative traces show longer free-running period in *Smg6*^*mut*^ cells (yellow) as compared to control cells (blue). **B**. Quantification of several experiments as in A. *Smg6* mutants (yellow) showed significantly longer periods in comparison to controls (*Smg6*^*flox/flox*^ treated with vehicle in blue, or *Smg6*^*+/+*^ with/without 4-OHT in grey). N=5 for *Smg6*^*flox/flox*^ cells; N=3 for *Smg6*^*wt/wt*^ cells; period difference between 4-OHT and vehicle-treated *Smg6*^*flox/flox*^ cells is 1.6 hours; Bonferroni’s multiple comparisons test adjusted p=0.0228. **C**. Adult *Smg6*^*flox/flox*^ male mice and their *Smg6*^*+/+*^ littermates (all carrying tamoxifen-activatable A*lbCreERT2* and the circadian reporter *Per2::Luc*) were sacrificed 4-5 weeks after tamoxifen injections. Liver and kidney explants were used for real-time recording of luciferase rhythms. Representative traces from livers show strong free-running period lengthening in the *Smg6* mutant (yellow) vs. control (blue). **D**. Quantification of several experiments as in C. Robustly increased periods, by almost 3 hours, were observed in NMD-deficient liver explants (yellow; mean=25.36 ± 2.23 h) in comparison to control livers (blue; mean = 22.0 ± 0.90 h). No period length difference was observed for kidney explants from the same animals (grey; flox/flox mean±SD = 25.2 ± 1.19 h, control mean±SD = 24.4 ± 1.83 h). Liver tissue: N=16 slices *Smg6*^*flox/flox*^; N=17 slices for controls; Mann-Whitney test p<0.0001. Kidney tissue: N=16 for *Smg6*^*flox/flox*^; N=20 for controls, Mann-Whitney test p=0.0771. 1-4 tissue slices were used per mouse; analyses performed blindly. **E**. Efficient recombination was validated by genotyping of livers (PCR analysis of genomic DNA). **F**. Cartoon depicting the *in vivo* recording setup (RT-Biolumicorder); *Smg6*^*flox/flox*^ or *Smg6*^*+/+*^ mice carrying *AlbCreERT2* and *mPer2::Luc* alleles were implanted with a luciferin-loaded mini osmotic pump 4 weeks after tamoxifen injections before placing in the recording device. **G**. Left panel: After 2 days under LD12:12, bioluminescence rhythms (photons) and activity (infrared signal) were recorded for 5 days under photoskeleton photoperiod conditions (indicated by yellow vertical lines at ZT12 and before ZT0). The plot shows mean signal (solid trace) and SEM (shaded) over the whole course of the experiment in the left panel, and compiled data, averaged from all days and mice, in the right panel. **H**. Quantification of PER2::LUC bioluminescence signal showed no difference in phase between tamoxifen-injected *Smg6*^*flox/flox*^ (yellow) and *Smg6*^*+/+*^ (blue) control animals. N=6 per group; Mann-Whitney test p=0.7251.

*In vivo*, and according to oscillator theory ^27,28^, a difference in period lengths between the entraining clock (here: wild-type period SCN) and the entrained clock (here: long period *Smg6* mutant hepatocytes) will typically translate to a phase shift of the entrained oscillator. Thus, we expected that the long period mPER2-Luc rhythms seen in liver explants *ex vivo* would lead to a change in phase *in vivo*. To evaluate this prediction, we used a method for the real-time recording of daily liver gene expression in freely moving mice ^29,30^ that relies on luciferase reporters, luciferin delivery via an osmotic minipump, and highly sensitive bioluminescence photon counting (**Fig. 2F**). Using the same *mPer2::Luc* reporter knock-in animals (NMD-deficient vs. controls) as for the above tissue explant experiments, real-time recording was carried out under conditions that ensured light-entrainment of the SCN clock to an external 24-hour light-dark cycle by means of a skeleton photoperiod, i.e. two 30 min light pulses applied at times corresponding to the beginning and to the end of the light phase in a 12h-light-12h-dark (LD12:12) cycle. We observed high-amplitude rhythmic bioluminescence rhythms in both genotypes (**Fig. 2G**) with phases that were, however, indistinguishable (**Fig. 2H)**. Next, we also investigated the effect of the *Smg6* mutation on the central clock in the SCN. We stereotactically injected an adeno-associated virus (AAV) expressing Cre::eGFP to induce recombination (**Fig. S1A, B**) and scored circadian clock parameters by two different assays: *in vivo*, we measured behavioral locomotor rhythms under constant conditions (free-running clock) by running wheel assays (**Fig. S1C, D**) and *ex vivo*, we recorded mPER2::LUC rhythms from SCN explants (**Fig. S1E, F**). Neither assay revealed an effect of the *Smg6* mutation on free-running periods for the SCN clock; yet, as a caveat, we also noted overall less efficient recombination as compared to our liver experiments (**Fig. S1G**). We concluded that loss of NMD triggered by the *Smg6* mutant allele had a strong period lengthening effect, notably for peripheral clocks and in particular in liver explants.

### NMD inactivation differentially affects the phases of core clock gene expression in the entrained liver

We next analyzed the apparent discrepancy between the long periods of liver rhythms *ex vivo* (**Fig. 2C, D**) and the lack of a phase phenotype *in vivo* (**Fig. 2G, H**). Briefly, other tissues than liver (e.g. kidney ^31^) may have contributed to the overall bioluminescence signal detected in the *in vivo* recording experiments, thereby masking a hepatic phase phenotype. Moreover, systemic cues that are dependent on the SCN, yet do not require a functional hepatocyte clock, can drive rhythmic PER2 accumulation in liver ^32,33^; therefore, mPER2::LUC signal may not be representative of the intrinsic liver clock phase. In order to evaluate in a comprehensive fashion how rhythmic gene expression was altered *in vivo*, we collected livers at 4-hour intervals around-the-clock from LD12:12-entrained *Smg6* mutant and control mice, with timepoints ZT0 (*Zeitgeber* Time 0, corresponding to time of “lights-on”), ZT4, ZT8, ZT12 (“lights-off”), ZT16 and ZT20 (**Fig. 3A**). We carried out RNA-seq on all individual mouse liver samples (triplicates per genotype and timepoint) and assembled the data into two time series representing the diurnal liver transcriptome under conditions of an inactive vs. active NMD pathway. As a means of quality control, we first validated that known NMD targets were upregulated in *Smg6* mutant livers. Indeed, as in the fibroblasts (**Fig. 1H, I**), NMD-annotated isoform exons were increased in abundance (**Fig. S2A**). Other transcripts diagnostic for an inactive NMD pathway showed the expected post-transcriptional upregulation as well. For example, mRNAs encoding components of the NMD machinery itself were post-transcriptionally upregulated (**Fig. S2B**), as reported previously from cell lines ^7^. This phenomenon has been proposed to represent an autoregulatory mechanism that involves as NMD-activating features the long 3’ UTRs that these mRNAs carry. Similarly, the uORF-regulated *Atf4* and *Atf5* transcripts, which are documented NMD substrates ^5,34^ and encode key transcription factors in the integrated stress response (ISR) ^35^, showed the expected upregulation (**Fig. S2C**). Of note, higher ATF5 protein accumulation (**Fig. S2D, E**) occurred in the absence of general ISR activation (as judged by eIF2α phosphorylation levels that were only weakly affected; **Fig. S2D**), pinpointing the lack of direct NMD regulation rather than proteotoxic stress as the likely trigger.

**Figure 3.**
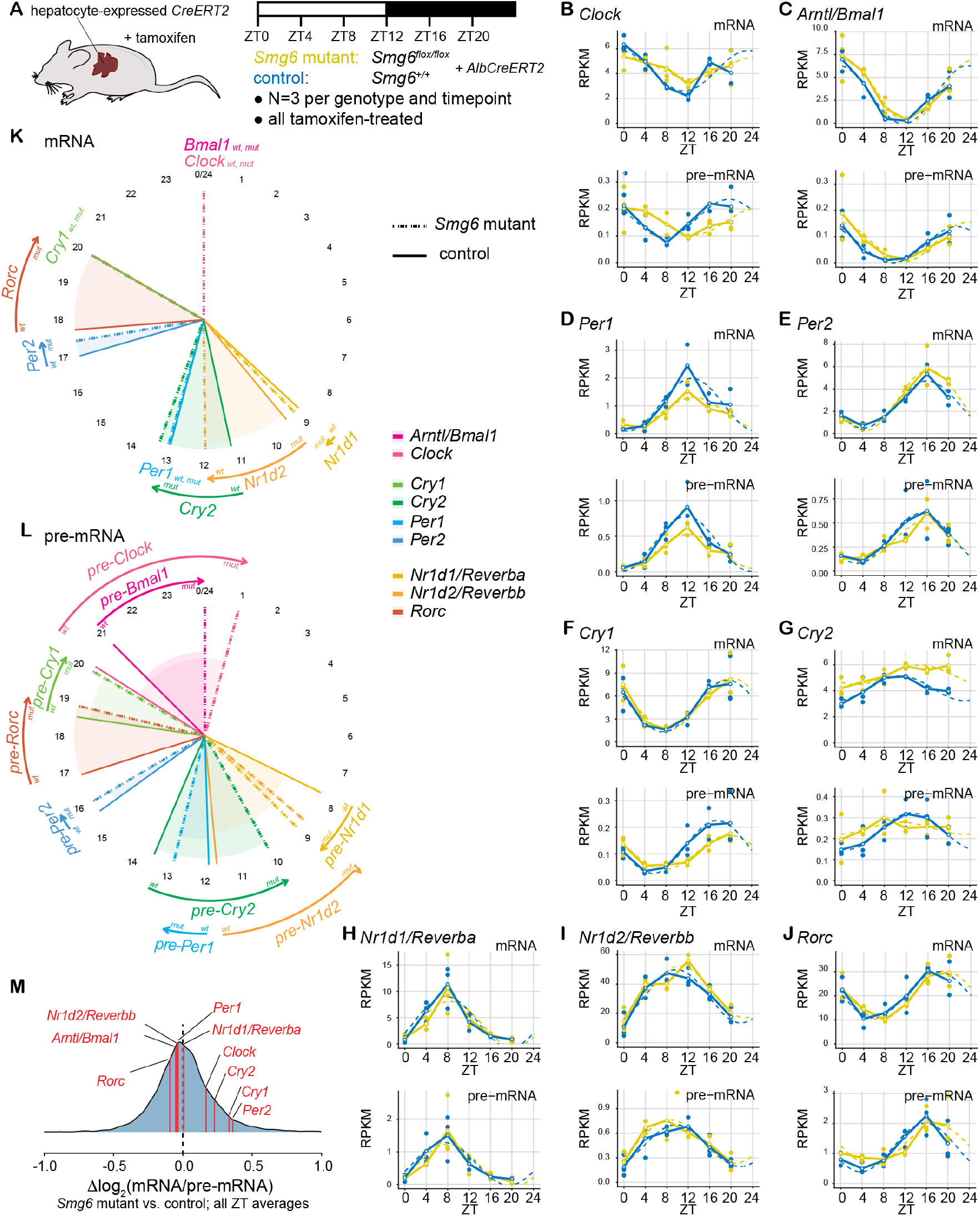
*Smg6* mutation differentially affects hepatic core clock pre-mRNA and mRNA rhythms. **A**. Schematic of the around-the-clock RNA-seq experiment, which was carried out on a time series of liver samples collected from LD12:12-entrained male *Smg6* mutant (*Smg6*^*flox/flox*^; *AlbCreERT2*; tamoxifen-treated) and control (*Smg6*^*+/+*^; *AlbCreERT2*; tamoxifen-treated) mice. **B.-J**. RNA-seq data is plotted for indicated core clock genes for mRNA (upper panels; exonic reads) and pre-mRNA (lower panels; intronic reads) for *Smg6* mutants (yellow) and controls (blue). RPKM values (Reads Per Kilobase of transcript, per Million mapped reads) of individual mice are shown as dots with solid lines connecting the means for each timepoint. The dashed lines represent the rhythmic data fit using the parameters from Metacycle ^36^. **K**. Circular plot representing the phases of peak mRNA abundances according to the Metacycle fits for *Smg6* mutants (dashed) and controls (solid) for indicated core clock genes. *Cry2, Nr1d2* and *Ror*γ accumulated several hours later in *Smg6* mutants, whereas minor effects were seen for the other genes. **L**. As in K. but for pre-mRNA rhythms. Several core clock pre-mRNAs showed later phases, indicative of transcriptional shifts; notable exceptions being *Per2* (almost invariable) and *Cry2* and *Nr1d2*, which both showed a phase advance. **M**. Similar to Fig. 1F, mRNA/pre-mRNA ratios were calculated for the liver RNA-seq data; briefly, average mRNA counts were first averaged over all samples per genotype, before dividing by average pre-mRNA counts. Three components of the negative limb, *Cry2, Cry1* and *Per2*, show higher mRNA/pre-mRNA ratios in *Smg6* mutants.

We then analyzed the daily dynamics of core clock gene expression at the mRNA and pre-mRNA levels (**Fig. 3B-J**). Consistent with the *in vivo* recording of *mPer2::Luc* animals, *Per2* mRNA and pre-mRNA rhythms were highly similar between the two genotypes (**Fig. 3E**). By contrast, several other core clock genes - notably those encoding the main transcriptional activators, *Clock* and *Arntl/Bmal1* (**Fig. 3B, C**), as well as *Cry1* (**Fig. 3F**) and *Rorc* (**Fig. 3J**) - showed phase-delayed pre-mRNAs indicative of transcription occurring several hours later. The complete analysis of core clock mRNA (**Fig. 3K**) and pre-mRNA (**Fig. 3L**) rhythms revealed that the considerable phase differences seen for many core clock genes at the transcriptional (pre-mRNA) level (**Fig. 3L**), only partially propagated to the mRNA level (**Fig. 3K**). Of the core loop constituents, *Cry2* mRNA showed a substantial delay by ca. 2 hours (**Fig. 3G, K**). The later phase did not have its origins in the timing of transcription, which rather appeared to be advanced (**Fig. 3G, L**). Other delays in mRNA accumulation that we observed affected the two nuclear receptors and components of the stabilizing loop, *Nr1d2/Rev-erbb* and *Rorc* (**Fig. 3H, J, K**).

### NMD regulation of *Cry2* mRNA occurs through its 3’ UTR and limits CRY2 protein accumulation in the dark phase

Among the core clock genes, the observed change in the daily *Cry2* expression profile (i.e. a peak in *Cry2* mRNA levels at ZT8-12 with subsequent decrease in control animals; yet *Cry2* mRNA abundances persisting on a high plateau until ZT20 in *Smg6* mutants; **Fig. 3G**) was consistent with the hypothesis that the *Cry2* transcript became stabilized in the absence of NMD. Indeed, the analysis of *Cry2* mRNA/pre-mRNA ratios across all liver samples suggested elevated stability during the dark phase of the cycle (ZT12-20) (**Fig. 4A**). Western blot analysis of total liver proteins revealed that the prolonged mRNA abundance under NMD-inactive conditions led to corresponding changes in the levels of CRY2 protein, whose peak accumulation was delayed by 4 hours in *Smg6* mutant animals (peak at ZT20) compared to controls (peak at ZT16) (**Fig. 4B, C**). Moreover, the analysis of individual livers showed that CRY2 reproducibly accumulated to >2x higher levels in *Smg6* mutant livers towards the end of the dark phase, at ZT20 (**Fig. 4D, E**). Furthermore, increased CRY2 levels were also apparent in *Smg6* mutant fibroblasts (**Fig. 4F, G**). These observations were consistent with a direct regulation of *Cry2* mRNA stability through NMD. To explore this hypothesis, we analyzed whether the *Cry2* mRNA contained any specific NMD-activating features. First, we inspected RNA-seq coverage on the *Cry2* locus in our fibroblast data, which revealed the expression of a single *Cry2* transcript isoform carrying a long 3’ UTR of ∼2.2 kb (**Fig. 4H**; identical observations were made in the liver RNA-seq data; data not shown), i.e. well beyond the ∼1 kb cut-off that has been used as a benchmark for the definition of potential endogenous NMD substrates ^2,6,7^. There was no evidence that a second annotated mRNA isoform with a shorter 3’ UTR (∼0.4 kb; **Fig. 4H**) or any other, additional transcript variants were generated from the locus. Finally, with a 5’ UTR that is particularly short (20 nt) and no evidence for translating ribosomes upstream of the annotated start codon according to previous ribosome profiling data from liver ^21^ or murine fibroblasts ^37^ (data not shown), we excluded the possibility that the transcript contained NMD-activating uORFs. We thus assessed whether the ∼2.2 kb *Cry2* 3’ UTR would confer NMD regulation to a luciferase reporter gene (**Fig. 4I**). Dual luciferase assays revealed that inactivating NMD in fibroblasts led to a >5-fold activity increase for the *Cry2* 3’ UTR-carrying reporter as compared to the control reporter (**Fig. 4J**), providing evidence that the *Cry2* 3’ UTR acts to elicit NMD.

**Figure 4.**
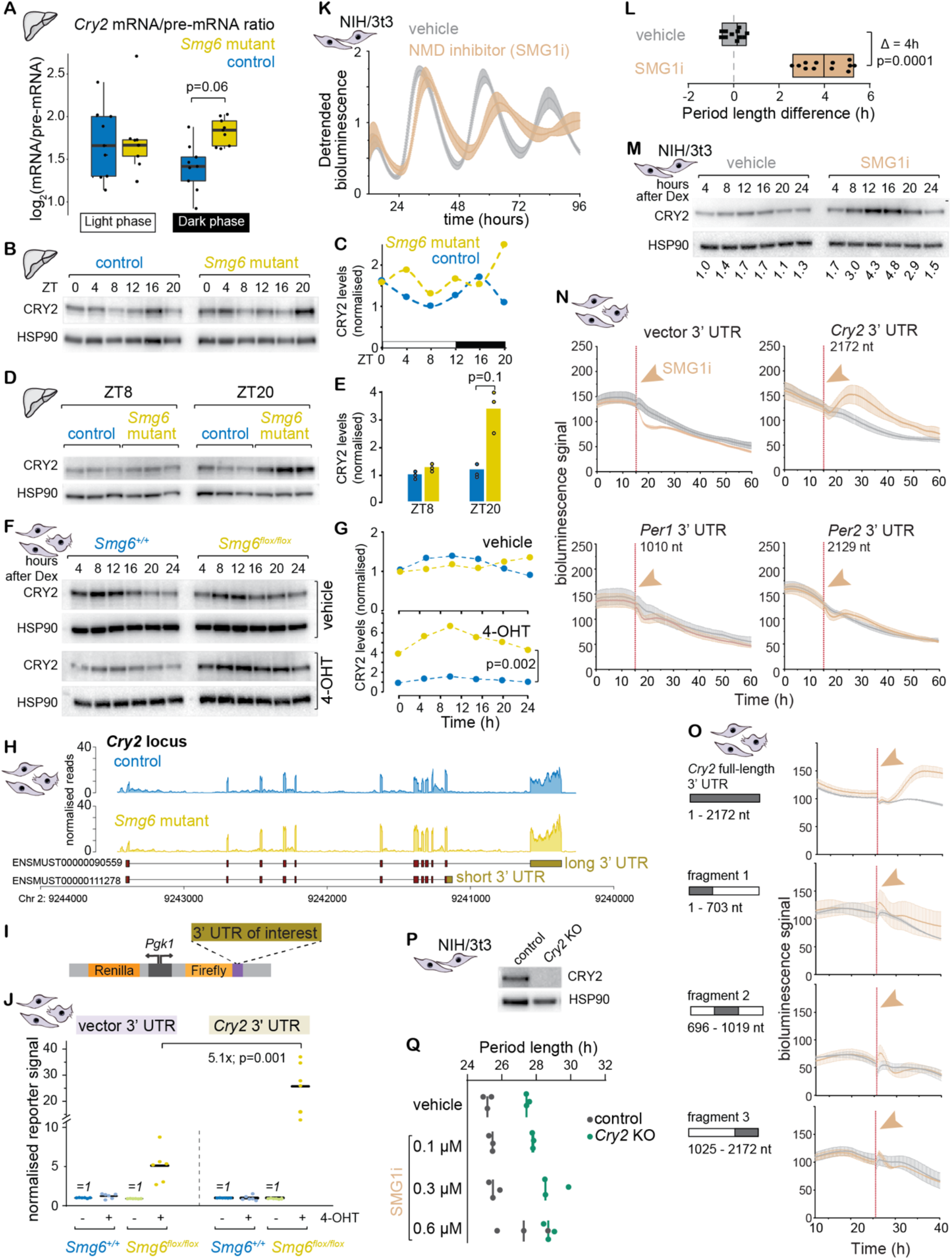
NMD regulation of *Cry2* mRNA via its 3’ UTR. **A**. mRNA/pre-mRNA ratios across individual liver samples – grouped into light (ZT0, 4, 8) and dark phase (ZT12,16, 20) samples – indicate *Cry2* mRNA stability increase in *Smg6* mutants, which is visible in particular in the dark phase; p-value=0.06; ANOVA. **B**. Western blot analysis of total liver proteins, for CRY2 and HSP90 (loading control). Each sample is a pool of 3 individual mice. **C**. Quantification of Western blot shown in B; CRY2 intensity was normalized to the loading control, HSP90. **D**. Western blot as in panel B, but from individual animals at ZT8 and ZT20, indicating that CRY2 is reproducibly more abundant at ZT20 in *Smg6* mutants. **E**. Quantification of Western blot in D; p=0.1; Mann-Whitney non-parametric test. **F**. Western blot analysis of total protein extract from fibroblasts (cells as shown in Fig. 1C) reveals CRY2 upregulation specifically in 4-OHT-treated *Smg6*^*flox/flox*^ cells. **G**. Quantification of Western blot shown in F; p=0.002; Mann-Whitney non-parametric test. **H**. RNA-seq read coverage on the *Cry2* locus (fibroblasts). Only one transcript isoform - carrying the 2.2 kb 3’ UTR - is expressed; there is no evidence for expression of the short UTR isoform. **I**. Schematic of the lentiviral dual luciferase system used in assays to test 3’ UTRs of interest for NMD regulation. **J**. Dual-luciferase assays reveal that the *Cry2* 3’ UTR confers NMD regulation. Non-treated cells for each genotype/reporter were internally set to 1. Vector UTR alone shows ca. 5-fold upregulation under *Smg6*^*flox/flox*^ + 4-OHT conditions (an effect coming from both Firefly luciferase up- and Renilla luciferase downregulation). Against this background of the assay, the *Cry2* 3’ UTR confers an additional >5-fold increase. Cells of each genotype/reporter condition without 4-OHT treatment were internally set to 1, and the signal of 4-OHT-treated cells relative to these untreated cells is reported; N=5 from 3 different experiments; p=0.001; Mann-Whitney non-parametric test. **K**. Bioluminescence traces of NIH/3t3 cells carrying the DBP-Luciferase reporter, with (orange) and without (grey) 0.6 µ M SMG1i treatment. Traces show average (mean) signal and standard deviation from 3 independent experiments. **L**. Quantification of experiments shown in K, showing reproducible period lengthening by ca. 4h in the presence of 0.6 µ M SMG1i (N=11-12; p<0.001; Mann-Whitney test). **M**. Western blot analysis of total protein extract from NIH/3t3 cells treated with vehicle or 0.6 µ M SMG1i; values of CRY2 abundance normalized to HSP90 (loading control) below the lanes. **N**. Primary fibroblast (genotype *Smg6*^*+/+*^, no 4-OHT) were stably transduced with luciferase reporters carrying different 3’ UTRs (as in panel I). Real-time recording of Firefly luciferase signal was carried until the signal reached stable state, before addition of 1 µ M SMG1i (orange) or vehicle (grey). The reporter carrying the *Cry2* 3’ UTR was specifically upregulated, as compared to *Per1, Per2* or vector 3’ UTRs. Traces show average (mean) signal and standard deviation; N=3. **O**. In assays as in N., only full-length *Cry2* 3’ UTR showed upregulation upon SMG1i treatment, but not individual fragments (N=2). **P**. Western blot showing absence of CRY2 in Crispr/Cas9-generated Cry2 knockout NIH/3t3 cells. **Q**. Period length of *Dbp*-luciferase reporter traces in NIH/3t3 cells - controls (grey) or *Cry2* knockouts (green) *-* treated with 0.1 µM, 0.3 µM or 0.6 µM of SMG1i or with vehicle (DMSO) corresponding to the volume used in highest SMG1i treatment.

We wished to further validate NMD regulation of the *Cry2* 3’ UTR by an approach that would allow more rapid and direct readout of reporter activity after NMD inhibition, rather than having to rely on prolonged 4-OHT treatment of reporter-expressing cells to induce the *Smg6* mutation. To this end, we used a pharmacological inhibitor of the kinase SMG1, hSMG-1 inhibitor 11e (SMG1i in the following) ^38^. Briefly, for this compound an IC50 in the sub-nanomolar range had originally been reported ^38^, yet subsequent studies *in vitro* ^39^ and in cells (e.g. ^40^) have applied SMG1i at considerable higher concentrations (0.2-1 μM) to inhibit NMD; additional effects on other kinases (e.g. mTOR ^38^) cannot be excluded under these conditions. Indeed, we observed a strong effect of 0.6 μM SMG1i on circadian period in two commonly used circadian model cell lines, murine NIH/3t3 fibroblasts (**Fig. 4K, L**) and human U2OS osteosarcoma cells (**Fig. S3**). Of note, the period lengthening phenotype caused by the compound (∼4 hours; **Fig. 4L**) was considerably stronger than that seen in the genetic *Smg6* fibroblast model (∼1.5 hours; **Fig. 2B**), in line with possibly broader activity of SMG1i. Moreover, cellular toxicity was observable after prolonged SMG1i treatment for several days (data not shown). We thus concluded that this compound would be most appropriate for short-term NMD inhibition up to 24 hours, which is also the timeframe in which it increased endogenous CRY2 protein abundance (**Fig. 4M**). We then assessed how acute SMG1i treatment affected the activity of lentivirally delivered luciferase reporters carrying various core clock gene 3’ UTRs, using real-time bioluminescence recording in mouse fibroblasts. Upon addition of SMG1i, output from a reporter carrying the *Cry2* 3’ UTR increased rapidly within a few hours (**Fig. 4N**). By contrast, neither the vector 3’ UTR, nor the 3’ UTRs of other core clock genes that were similar in length to the *Cry2* 3’ UTR, namely that of *Per1* (∼1 kb) and *Per2* (∼2.1 kb), showed increased reporter output. Based on this outcome, we concluded that the *Cry2* 3’ UTR was a specific target of the NMD pathway. We next reasoned that the *Cry2* 3’ UTR may be NMD-activating due to its length or, alternatively, that it could contain specific *cis*-acting elements important for NMD activity, e.g. specific binding sites for RNA binding proteins (RBPs). To distinguish between these two scenarios, we tested individual, overlapping fragments of the full-length *Cry2* 3’ UTR in the reporter assay. In contrast to full-length *Cry2* 3’ UTR, none of the fragments was associated with reporter upregulation upon SMG1i treatment (**Fig. 4O**). We concluded that most likely the considerable length of the *Cry2* 3’ UTR was responsible for downregulation via NMD.

With NMD downregulation leading, on the one hand, to longer periods and, on the other hand, to altered abundance and accumulation dynamics of CRY2, we next attempted to investigate whether there was a causal link between both effects. To this end, we produced *Cry2*-deficient NIH/3t3 cells (**Fig. 4P**). We treated these cells with SMG1i, based on the reasoning that NMD inhibition may have a less severe phenotype in the absence of a functional *Cry2* gene. However, in this setup we did not uncover an evident modulation of SMG1i-mediated period lengthening by the absence of *Cry2* (**Fig. 4Q**). A similar outcome was obtained in *Cry2*-deficient U2OS cells (**Fig. S3**). We concluded that the SMG1i-provoked period phenotype was not dependent on *Cry2*. However, given the questions surrounding the specificity of SMG1i detailed above, an interaction of the phenotype with *Cry2* may have been masked by other, stronger effects of the compound. Dedicated experiments using *Smg6*^*mut*^ cells/livers will thus be required in the future to evaluate to what extent NMD-mediated regulation of *Cry2* contributes to period lengthening.

### Transcriptome-wide analyses uncover the extent of rhythmic gene expression reprogramming in the entrained liver

We next analyzed how, beyond the core clock genes (**Fig. 3**), the global rhythmic transcriptome was affected in *Smg6* mutant livers *in vivo*. Our expectation was that we would find a complex overlay of direct and indirect effects, due to (i) NMD directly controlling the mRNA stability for some clock-controlled output genes, which would post-transcriptionally impact on their amplitudes and phases; (ii) the altered phase of *Cry2* and other core clock components (**Fig. 3K**) impacting on the transcriptional timing and dynamics at clock-controlled loci; and (iii) additional secondary consequences that could be both transcriptional and post-transcriptional in nature, as a result of the above effects. We first investigated whether there were global changes in the populations of rhythmic transcripts between the two genotypes, analyzing the RNA-seq datasets from the above cohort (**Fig. 3A**). Using established rhythmicity detection algorithms (MetaCycle R package ^36^), we found that the majority of mRNAs classified as rhythmic in controls were also rhythmic in the *Smg6* mutant livers (N=1257; **Fig. 5A**) and visual inspection of the pre-mRNA heatmaps further suggested that most of these rhythms were of transcriptional origin. A lower number of transcripts passed the rhythmicity criteria in only one of the genotypes, indicating possible loss (N=223; **Fig. 5B**) or gain (N=323; **Fig. 5C**) of oscillations in the *Smg6* mutants. Inspection of the heatmaps, however, indicated that in many cases, the alleged lack of rhythmicity in one or the other genotype was probably the result of effects such as lower/noisier expression levels rather than clear-cut loss of daily oscillations (a well-known phenomenon when comparing rhythmic gene expression datasets, see ^41,42^). We thus first focused our analyses on the common mRNA rhythmic genes. Their peak phase distributions globally resembled each other in the two genotypes (**Fig. 5D**). A large group of mRNAs showed maximal abundance around ZT6-12 (an interval that overlaps with the expected peak mRNA phase of direct BMAL1:CLOCK targets containing E-box enhancers ^43^), and this cluster appeared phase-advanced in *Smg6* mutants. Moreover, several phases were underrepresented in mutants as compared to controls, such as the distinct group of transcripts with maximal abundance at the beginning of the light phase (ZT0-2) in controls that was absent in *Smg6* mutant livers (**Fig. 5D**). For a more quantitative analysis of these effects, we calculated transcript-specific phase differences, which indicated that mRNA phases in *Smg6* mutants globally followed those in controls, with advances and delays spread out across the day (**Fig. 5E**). Overall, more transcripts were phase advanced than delayed in *Smg6* mutant livers (**Fig. 5F**). This outcome was unexpected given that the expression profiles for core clock transcripts (**Fig. 3B-J**), and specifically the findings on *Cry2* (**Fig. 4**), had rather pointed towards a delay of the entrained liver clock in *Smg6* mutants. To investigate these observations further, we overlaid our rhythmic transcript set with data from a large circadian mouse liver ChIP-seq study ^20^. Our analyses revealed that mRNAs arising from loci with binding sites for BMAL1 and CLOCK (**Fig. 5G**) or PER and CRY proteins (**Fig. 5H**) were indeed significantly skewed towards phase delays, in contrast to rhythmic genes that were not direct targets of these core clock proteins (**Fig. 5I**). We concluded that multiple factors engendered phase changes at the rhythmic transcriptome level in *Smg6* mutants, manifesting in delays for many direct BMAL1:CLOCK targets, and overall advanced phases for many other rhythmically expressed mRNAs. Next, we compared peak-to-trough amplitudes between the genotypes, given that for rhythmic mRNAs that are direct targets of NMD, increased transcript stability in *Smg6* mutants should lead to amplitude reduction. To explore this possibility, we used the Z-scores (**Fig. 5A**) for the common rhythmic transcripts to calculate the amplitudes (maximum-to-minimum fold-changes) for mRNAs and for pre-mRNAs, which we compared between the two genotypes. In *Smg6* mutants, median mRNA amplitudes were lower than in controls, but pre-mRNA amplitudes were higher (**Fig. 5J**); when normalizing mRNA amplitudes for pre-mRNA fold-changes – as a means to control for differences in transcriptional rhythmicity at the locus – the decrease in rhythmic transcript amplitudes in *Smg6* mutants became highly significant (**Fig. 5K**). This outcome indicated that higher stability of rhythmic mRNAs in *Smg6* mutants was detectable at the global level. In the extreme scenario, an mRNA that is rhythmic in control animals would lose its amplitude to the extent that it would not anymore be considered as rhythmic at all; it would then group within the N=223 genes shown in **Fig. 5B**. We inspected their individual gene expression profiles, which led to the identification of a sizeable number of transcripts that displayed severely blunted mRNA amplitudes in *Smg6* mutants, despite similar rhythmic pre-mRNAs (i.e. oscillations in transcription) (**Fig. 5L**). For several of the cases, we can speculate about possible NMD-eliciting features. For example, according to our previous mouse liver ribosome profiling data ^21^, *Glycine decarboxylase (Gldc)* contains translated uORFs (data not shown); in the case of *Lactate dehydrogenase B (Ldhb)*, a regulatory mechanism entailing stop codon readthrough has been demonstrated [31] and could potentially link *Ldhb* translation to NMD regulation. For the other transcripts shown in **Fig. 5L** (*Pde9a, Kyat1, Tubb4, Tmem101, Amdhd1, Epha2*), no obvious candidate NMD-eliciting features were found.

**Figure 5.**
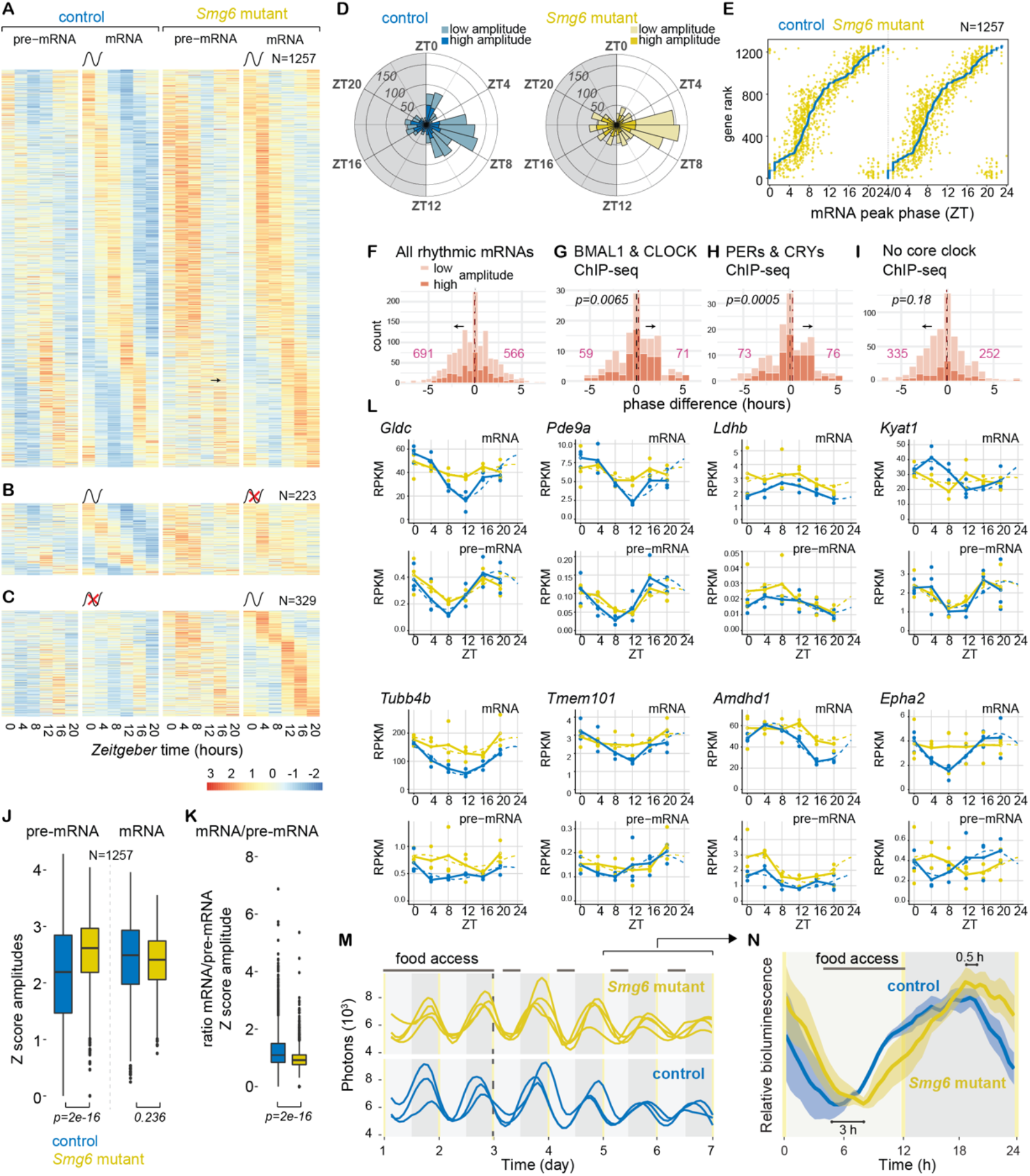
Rhythmic RNA expression is altered in *Smg6* mutant livers under entrainment. **A**. Heatmap of transcripts with significant rhythms at the mRNA level in both genotypes. Expression levels are represented as Z-scores for mRNA or pre-mRNA with color code for low (blue) to high (red) expression. Z-scores were calculated separately for mRNA and pre-mRNA data, but on a common scale for both genotypes. Transcripts are phase-order for the control genotype. **B**. Heatmap as in A., but for transcripts with significant rhythms only in control animals (N=223) and not in mutants. **C**. Heatmap as in A., but for transcripts with significant rhythms only in mutant animals (N=329) and not in controls. **D**. Radial diagrams showing peak phase of rhythmic mRNAs in control (blue) and *Smg6* mutant (yellow) liver for the common transcripts shown in A. Dark shaded: high amplitude rhythmic transcripts; light shaded: low amplitude rhythmic transcripts; high/low cut-off on log2 peak-trough amplitude of 1. **E**. Peak phase of mRNA in *Smg6* mutants (orange) relative to control phase (blue), ranked according to the phase in the control (*Smg6* wt) for the transcripts shown in A (N=1257). **F**. Peak phase difference between mutant and control mice for all common rhythmic mRNAs (N=1257). **G**. Peak phase difference of commonly rhythmic mRNAs as in F., but restricted to loci with ChIP-seq binding sites for BMAL1 and CLOCK (N=130), according to ^20^; p=0.0065; permutation test, calculated by 1000x subsampling of N=130 transcripts from the “all rhythmic transcripts” (N=1257) of panel F, then comparing the means of these subsampling groups with the observed mean (t-test). **H**. Peak phase difference of commonly rhythmic mRNAs as in F., but restricted to loci with ChIP-seq binding sites for the ensemble of proteins PER1, PER2, CRY1 and CRY2 according to ^20^; p=0.0005; permutation test as in panel G. **I**. Peak phase difference of commonly rhythmic mRNAs as in F., but restricted to loci with no ChIP-seq binding sites for any of the proteins BMAL1, CLOCK, PER1, PER2, CRY1 or CRY2 according to ^20^; p=0.18; permutation test as in panel G. **J**. Z-score amplitudes – defined as the difference between the maximum and minimum Z-score values, calculated independently for mRNAs and pre-mRNAs of commonly rhythmic transcripts (N=1257) - show lower mean mRNA (p=0.236) and higher mean pre-mRNA amplitudes in mutants (p=2e-16); significance calculations from a linear model (equivalent to t-test). **K**. Transcript mRNA/pre-mRNA Z-score amplitude ratios (from the N=1257 common rhythmic transcripts) stratified by genotype show decrease in mutants; p=2e-16; Student’s t-test. **L**. RNA-seq data is plotted for indicated genes for mRNA (upper panels; exonic reads) and pre-mRNA (lower panels; intronic reads) for *Smg6* mutants (yellow) and controls (blue). Rhythmicity of mRNA levels observed in control (blue) is dampened or lost in *Smg6* mutant liver (yellow). RPKM values of individual mice are shown as dots with solid lines connecting the means for each timepoint. The dashed lines represent the rhythmic data fit using the parameters from Metacycle ^36^. **M**. RT-Biolumicorder traces of individual mice in food shifting experiment. After 2 days under *ad libitum* feeding, bioluminescence rhythms (photons) and activity (infrared signal) were recorded for 4 additional days under light-phase-restricted feeding conditions (ZT10-20; horizontal black bar at top); skeleton photoperiod entrainment indicated by yellow vertical lines at ZT12 and ZT0. Each line represents the signal from a control (blue) or a liver-specific *Smg6* mutant (yellow) animal **N**. Compiled data, averaged over the last two days of the experiment. Mean signal (solid trace) and SEM (shaded). Indicated phase differences calculated from rhythmic fits to the data.

Collectively, these analyses demonstrated that the stably entrained liver clock, under *ad libitum* feeding and LD12:12 conditions, was subject to phase and amplitude alterations at the level of clock-controlled gene expression. Our *in vivo* recording experiments (**Fig. 2F-H**) had been insensitive to picking up such differences in liver rhythms due to the use of the *mPer2::Luc* reporter allele, whose phase was unaffected by *Smg6* mutation under stable entrainment conditions. We reasoned that under conditions where the stable entrainment was challenged, a phenotype may be unmasked also for *mPer2::Luc*. To this end, we carried out food shifting experiments i.e., switching from *ad libitum* to daytime feeding. Under these conditions, the liver clock receives conflicting timing cues from the SCN and from feeding/fasting cycles, which are not anymore aligned and will eventually lead to an inversion of hepatic oscillator phase due to the dominance of feeding signals for peripheral oscillators ^44^. The kinetics and endpoint of phase adaptation can also be understood as a paradigm of clock flexibility and can be recorded using the RT-Biolumicorder setup ^30^. Our experiments showed that in *Smg6* mutant animals, after 3 days of feeding during the light phase, daily cycles in bioluminescence had readjusted to a new phase that substantially differed between control and *Smg6* mutant animals (3 hours difference at trough/0.5 hours at peak; **Fig. 5M, N**). We concluded that NMD contributes to the adaptation of circadian gene expression to food entrainment in mouse liver. More generally, the data point to notable differences between *Smg6* mutant and control animals with regard to how different timing cues are integrated within the core clock circuitry.

## Discussion

Our novel conditional *Smg6* endonuclease-mutant allele provides unique possibilities to explore *in vivo* activities of the NMD pathway and has allowed us to uncover an unexpected role within the mammalian circadian system, which is a conserved, key mechanism for the organization of daily rhythms in behavior, physiology and metabolism. We find that NMD loss-of-function has a striking impact on free-running circadian periods in two peripheral clock models, primary fibroblasts and liver. Moreover, we determine a specific core clock component, *Cry2*, as NMD-regulated and attribute the NMD-eliciting activity to its long 3’ UTR. Although it is widely accepted that efficient mRNA decay is critical for the establishment of gene expression oscillations, which specific pathways mediate the decay of transcripts encoding core clock components has remained largely unknown. That NMD has been co-opted for this purpose, as we find to be the case for *Cry2*, is surprising at first sight – yet it may simply reflect that nature and evolution are opportunistic and employ the available molecular pathways in the most efficient fashion. In line with this idea is the finding that a sizeable number of other rhythmic transcripts appears to rely on NMD to ensure efficient mRNA turnover as well (**Fig. 5L**). Our observations may change the way we should perceive the evolutionary drives relating to NMD: for example, it has been speculated why many mammalian mRNAs contain long 3’ UTRs but evade NMD, and a model has been put forward suggesting that such mRNAs have evolved to recruit NMD-inhibiting RBPs in spatial proximity of the termination codon ^6^. However, an opposite drive to attract and retain NMD regulation would be plausible as well – acting on endogenous transcripts, such as *Cry2*, whose intrinsic instability is physiologically important. This idea is in line with findings that in the circadian systems of *Neurospora* ^15^, *Arabidopsis* ^13^ and *Drosophila* ^14^, roles for NMD have emerged as well.

In the absence of NMD, CRY2 protein in liver accumulates to higher levels and for an extended time (**Figure 6**). Based on the experiments presented in our study, we are not yet fully in the position to evaluate to what extent these effects are involved in the period lengthening phenotype. Still, it would be plausible that the phase delay of CRY2 seen in the *Smg6* mutants could be particularly critical. According to around-the-clock ChIP-Seq data from wild-type mouse liver, CRY2 binds and represses its target genes at circadian time CT15-16 ^20^, thus closely matching the timing of maximal CRY2 abundance in our control mice (ZT16). The ChIP-seq data from wild-type livers further indicates that by CT20, CRY2 is cleared and replaced by CRY1, which binds to chromatin with a peak at around CT0 and is associated with a transcriptionally repressed, but poised state of BMAL1:CLOCK activity. Period lengthening through the prolonged availability of CRY2 may thus involve an extended CRY2-mediated repressive phase and/or CRY2 denying its homolog CRY1 access to its targets, causing a delay in the handover to CRY1. Of note, the period lengthening we observe is phenotypically comparable to that reported for a chemical, selective stabilizer of CRY2 protein, which also prolongs period in reporter assays across several cell types and species ^45^. Moreover, period lengthening has also been reported upon CRY2 stabilisation (in a *Cry1*-deficient background) induced by genetic inactivation of the CRY-specific ubiquitin ligase *Fbxl3* ^46^. For these reasons – and reminiscent of findings on CRY1 accumulation ^47^ – the changed timing of CRY2 accumulation, rather than its generally higher levels, may be a critical feature for the period phenotype and for the phase effects seen in the entrained liver. We thus propose that limiting temporal *Cry2* mRNA availability, mediated through NMD, is an important mechanism within the core loop of the clock by which CRY2 protein biosynthesis is restricted to the beginning of the dark phase when it acts in sync with PER1 and PER2 to repress CLOCK:BMAL1-mediated transcription. Only after duly removal of this repressive complex can CRY1 join and advance the cycle through the late repressive and poised states, eventually leading to the next transcriptional cycle at CLOCK:BMAL1-bound E-box enhancers.

**Figure 6.**
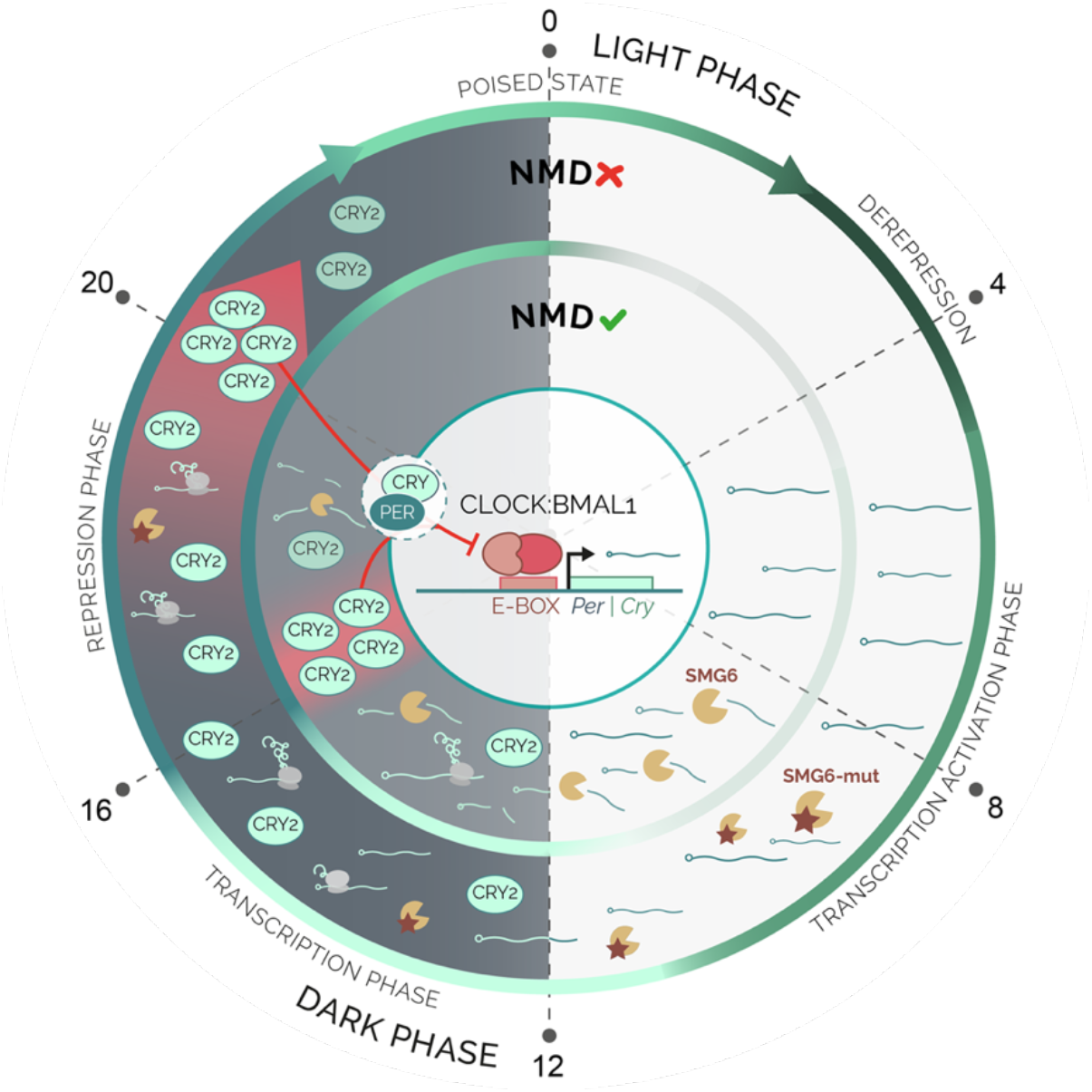
Model of how the daily dynamics of CRY2 accumulation are regulated by NMD. In the entrained liver clock, *Cry2* mRNA is translated and the protein accumulates with a peak in the dark phase (ZT16 in wild-type). In the absence of a functional NMD pathway, *Cry2* mRNA is stabilized, reaches higher levels, and its translation leads to increased CRY2 at later times (ZT20). The specific phases and states noted at the periphery of the circle (poised, derepression etc.) refer to the findings from Koike *et al*. ^20^ on E-box binding of clock proteins.

Intriguingly, our findings suggest specificity of the phenotype for peripheral clocks. Thus, we were unable to detect an impact on circadian period of the master clock in the SCN. Different explanations may underlie this observation. First, we cannot exclude lack of phenotype due to technical reasons, in particular the lower efficiency of Cre-mediated recombination in SCN neurons, or slow replacement kinetics of wild-type SMG6 by its mutant version due to high protein stability in neurons. For possible biological explanations, the decay of NMD substrates may be less reliant on SMG6 in neuronal cells, or the strong intercellular coupling in the SCN ^48^ renders the clocks resilient against the genetic NMD perturbation and the resulting changes in the critical NMD-regulated transcript. Finally, if the phenotype actually does involve CRY2, it is interesting that is has been reported that the relative importance of the two homologs, CRY1 and CRY2, in the negative feedback loop can be rather tissue-specific, with CRY1 being the main transcriptional repressor in the SCN ^46^, leading to another potential explanation for the observed cell type-specificity. Future experiments will be required to distinguish between these possibilities.

In summary, the unexpected role of NMD that we uncover within the circadian system illustrates the ongoing shift in perception of NMD from surveillance to housekeeping functions. We anticipate that our mouse model will provide valuable insights into so-far unidentified NMD targets and functions in mammals *in vivo*, including in the context of pathologies such as neurological diseases ^49^ and cancer ^50,51^, and in situations where NMD has been identified as a promising therapeutic target ^52,53^.

## Methods

### Animals

All animal experiments were performed according to the cantonal guidelines of the Canton of Vaud, Switzerland, license VD3611. Healthy adult male mice of age 12 – 24 months were used. All mouse lines were maintained on a C57BL/6J background. The alleles *AlbCre-ERT2*^*ki* 25^ and *mPer2::Luc*^*ki* 26^ have been previously described. The novel *Smg6*^*flox*^ allele was generated in collaboration with Taconic (official nomenclature of line: *Smg6tm5498(D1352A,D1391A)Tac*).

### Primary fibroblasts and immortalization

Adult male *Smg6*^*flox/flox*^ and *Smg6*^*+/+*^ control littermate mice were euthanized and approximately 1 cm of tail tip was recovered and further sliced into thin pieces under sterile conditions. Tissue fragments were overnight digested with 1 mg/ml collagenase type 1A (Sigma Aldrich) in culture medium at 37°C. The culture medium consists of 15% of fetal calf serum (FCS), 1% Penicillin-Streptomycin-Glutamine (Thermo Fisher Scientific, 10378016), 1% non-essential amino acids (Thermo Fisher Scientific, 11140050), 1 mM sodium pyruvate (Thermo Fisher Scientific, 11360070), 87 mM β-mercaptoethanol, 18 mM HEPES pH 7.0 (Thermo Fisher Scientific, 15630080), 2.5 µ g/ml Amphotericin B (Thermo Fisher Scientific, 15290018) and 2.5 µ g/ml Plasmocin (InvivoGen).

Isolated fibroblasts became spontaneously immortal upon continuous culture, creating *Smg6*^*flox/flox*^ or *Smg6*^*+/+*^ cell lines. Immortalized fibroblasts were transduced with a retrovirus carrying a tamoxifen-inducible Cre and puromycin resistance (MSCV *CreERT2* puro, Addgene plasmid #22776) ^54^. Retrovirus production was performed using the pCL-eco (Addgene, 12371) ^55^ and pCMV-VSV-G (Addgene, 8454) ^56^ plasmids in 293FT HEK cells using the CalPhos™ Mammalian Transfection Kit (Takara bio, 631312). Following 2 μM tamoxifen treatment, renewed every 24h for 4 consecutive days, the cells were utilized for experiments after 7-10 days from the treatment initiation.

### DNA genotyping

DNA from cell cultures, liver or kidney tissue was extracted using the DNeasy® Blood & Tissue Kit (Qiagen, 69504) according to the manufacturer’s protocol. Genotyping PCR reaction was performed using HotStar Taq DNA polymerase (Qiagen, 203207), 0.4 uM primers (Microsynth), 0.2 uM dNTP mix (PROMEGA, U1511) and approximately 200-700 ng of DNA template. The primer sequences are as follows (5’-3’): Forward: gaa ata cca ggg ccc ttg c, Reverse1: cat cac tac cca gct cag gaa c, Reverse2: gga ttg gct cct ctt tgc tg. The PCR program is as follows : 15 sec at 95°C, 35 cycles : 1 min at 94°C, 1 min at 61°C, 1 min at 72°C and final elongation at 72°C for 10 min. DNA extraction from dissected SCN tissue was done by Arcturus® PicoPure® DNA Extraction Kit (Thermo Fisher Scientific, KIT0103). PCR reaction was set up as above. The primer sequences are as follows (5’-3’): Forward gaa ata cca ggg ccc ttg c, Reverse2: tct agc tcc ttt ctg cct ctt c. The PCR program is as follows : 15 sec at 95°C, 40 cycles : 1 min at 94°C, 1 min at 55°C, 1 min at 72°C and final elongation at 72°C for 10 min.

### Luciferase reporters and lentiviral production

*CreERT2 Smg6*^*flox/flox*^ and *Smg6*^*+/+*^ immortalized fibroblasts were transduced with a lentivirus carrying a dual luciferase (Firefly/Renilla) NMD reporter or a control vector. For the generation of dual luciferase reporter plasmids, the prLV1 dual luciferase reporter plasmid ^11^ was used, with or without the introduction of an intron downstream of the *Firefly* stop codon. For the latter, the chimeric intron of the pCI-neo vector (Promega, E1841) was cloned into the 3’ UTR of the prLV1 vector. The following primers were used for PCR amplification: forward: aaagcggccGCTCGTTTAGTGAACCGTC (introducing a NotI restriction site) and reverse: tTTCTCGAGCTGTAATTGAACTGGGAG (introducing a XhoI restriction site). *Dbp-Luciferase* ^23^ and the 3’ UTR luciferase reporters ^11^ have been described previously. Lentiviral particles were produced in 293T cells using the envelope vector pMD2.G and the packaging plasmid psPAX2 as previously described ^57^. Filtered viral supernatant was spun 2h at 24,000 rpm, 4°C using Optima L-90K Ultracentrifuge (SW32Tirotor; Beckman, Brea, CA), then viral particles were resuspended with normal growth medium and used for cell transduction.

### Circadian bioluminescence recording of cell cultures

Fibroblasts cultured in 35 mm culture dishes (Falcon, 353001) were synchronized either with serum shock (50% horse serum for 3h) or with temperature entrainment (cycles of 16h at 35° C and 8h at 37°C for 5 days). During recording cells were cultured in phenol-free DMEM (Gibco, Thermo Fisher Scientific, 11880028) containing 10% FBS, 1% PSG and 0.1 mM of luciferin, sealed with parafilm to avoid evaporation, in the LumiCycler setup (Actimetrics) at 37°C and 5% CO2. NIH/3T3 murine fibroblasts were cultured under the same conditions as the immortalized fibroblasts but synchronized with 100 nM Dexamethasone treatment for 15 min. SMG1 inhibitor (hSMG-1 inhibitor 11e; Probechem Cat. No. PC-35788) ^38^ was used as 10 mM stock (dissolved in DMSO) and, if not indicated otherwise, used at a concentration of 0.6 μM (NIH/3T3 experiments) to 1μM (*Smg6*^*flox*^ fibroblasts).

### Dual Luciferase assay

After lentiviral transduction cells were collected using 5x Passive Lysis Buffer (Promega) and luciferase activity was measured using the Dual-Glo Luciferase Assay System (Promega, E1910) according to the manufacturer’s protocol. *Firefly-*Luciferase signal was normalized to *Renilla-*Luciferase, and for each construct (3’ UTR or NMD reporter) this signal was then normalized to that of lentivector-control plasmid (only containing generic vector 3′ UTR) treated with vehicle (for each experiment).

### RNA sequencing and analysis

Reads were mapped on the mouse genome GRCm38 (Ensembl version 91) using STAR ^58^ (v. 2.7.0f; options: --outFilterType BySJout --outFilterMultimapNmax 20 -- outMultimapperOrder Random --alignSJoverhangMin 8 --alignSJDBoverhangMin 1 -- outFilterMismatchNmax 999 --alignIntronMin 20 --alignIntronMax 1000000 -- alignMatesGapMax 1000000). Read counts in genes loci were evaluated with htseq-count ^59^ (v. 0.13.5) for transcript mapped reads (i.e. exons; options: --stranded=reverse --order=name --type=exon --idattr=gene_id --mode=intersection-strict) and for whole locus mapped reads (i.e. exons plus introns; options: --stranded=reverse --order=name --type=gene -- idattr=gene_id --mode=union). Read counting for exon analysis was not possible with htseq-count (most reads spanned multiple exons and would have been discarded) so a new python script was developed for this task. To avoid counting reads spanning different exons multiple times, the script calculated average read depth for each exon. Read pileups for gene loci were calculated using samtools depth ^60^ (v. 1.9) and plotted using R (v 4.1.1). Differential expression analysis was done in R using DESqe2 package ^61^. RNA stability analysis was performed using RPKM normalised reads counts. Phase analysis was performed using RPKM normalised reads counts and the MetaCycle R package ^36^.

### Induction of liver-specific *Smg6* mutation

8-12 week old male *Smg6*^*flox/flox*^ mice, carrying the liver-specific Albumin-driven CreERT2 (allele *Alb*^*tm1(cre/ERT2)Mtz* 25^), and their control littermates (*Smg6*^*+/+*^) received 4 intraperitoneal injections of 20 mg/ml tamoxifen (Sigma-Aldrich) in corn oil at a dosage of 75 mg tamoxifen/kg of body weight. The mice were admitted for experiments 4 weeks later.

### Liver and kidney explants

Male *Smg6*^*flox/flox*^ mice and their control littermates *Smg6*^*+/+*^ were euthanized following deep anesthesia by isoflurane inhalation. Liver and kidney tissue were excised and put immediately in ice-cold Hank’s buffer (Thermo Fisher Scientific). The outermost edges of the tissues were carefully excised in a sterile cabinet, and immediately placed on a 0.4 micron Millicell cell culture inserts (PICMORG50) in a 35 mm dish with phenol-free DMEM (Thermo Fisher Scientific, 11880028) containing 5% FBS, 2 mM glutamine, 100 U/ml penicillin, 100 μg/ml streptomycin and 0.1 mM luciferin. The parafilm-sealed plates were placed for recording in the LumiCycler (Actimetrics) at 37°C and 5% CO2.

### RT-Biolumicorder experiments

Adult male mice, 12-20 weeks of age, carrying the genetically encoded circadian reporter allele *mPer2::Luc* ^26^ were used for the RT-Biolumicorder experiments. The experimental procedure followed our recently published protocol ^29^. Briefly, Alzet mini-osmotic pumps (model 100D5 or 2001) were filled with 90 mg/ml with D-Luciferin sodium salt, dissolved in Phosphate Buffered Saline (PBS, pH 7.4) under sterile conditions. The pumps were closed with blue-colored flow moderators (ALZET) and activated at 37°C according to the manufacturer’s instructions, followed by the subcutaneous, dorsal implantation. As analgesics Carprofen (Rimadyl, 5 mg/kg subcutaneous), and paracetamol (2 mg/ml, via drinking water) were administered. Prior implantation the dorsal area of the mouse at the site where the liver is positioned was shaved using an electric razor. The RT-Biolumicorder (Lesa-Technology) consists of a cylindrical cage for a single mouse with photon-reflecting walls, equipped with a photomultiplier tube (PMT), water and food containers and a built-in infrared sensor that records locomotor activity (^29,30^). The RT-Biolumicorder records photon and activity levels in 1 min intervals. The data, which also contain light and food access information, were saved as text files and later analyzed using the MatLab-based “Osiris” software according to ^29^ or a custom-made R script.

### Running wheel experiments

12-16 week old male mice were single-housed in cages equipped with a running wheel and were placed in a light-tight cabinet. After approximately 10 days of habituation in 12h light-12h dark the mice were released in constant darkness for approximately 14 days. For the running wheel experiments with SCN-specific *Smg6* mutant recombination, the same protocol was used, followed by 14 days of post-injection recovery under 12h-light-12h-dark conditions and a second period of constant darkness for 14 days (adapted from ^62^).

### SCN-specific *Smg6* mutant mice

Male adult *Smg6*^*flox/flox*^ mice and their control littermates (*Smg6*^*+/+*^) received bilateral stereotactic injections of CMV.HI-Cre::eGFP AAV5 particles (AddGene, 105545) into the SCN (400 nl per site). Stereotactic coordinates: AP= - 0.34 ML= +/- 0.4, V=5.5. Ketamine/Xylazine (80/12.5 mg/kg) by intraperitoneal injection was used as anesthetic and 5 mg/kg carprofen was administered subcutaneously for analgesia. Additionally, paracetamol (2 mg/ml) was administered via drinking water prior and 3 days following the procedure. Animal recovery was monitored for ten days. Mice carrying *mPer2::Luc* ^26^ in addition to *Smg6*^*flox/flox*^ (experimental) or *Smg6*^*+/+*^ (control) were used for the bioluminescence recording of SCN slices. For evaluation of viral targeting, mice were transcardially perfused with phosphate-buffered saline (PBS) followed by 4% paraformaldehyde (PFA). Brains were post fixed overnight in 4% PFA at 4°C and then cryopreserved in 30% sucrose solution in PBS for at least 24 hours at 4°C (until completely sunk to the bottom of the container). Cryopreserved brains were frozen and sliced in 25 μm thick sections. Sections were mounted using DAPI-fluoromount. Fluorescent images were acquired on a ZEISS Axio Imager.M2 microscope, equipped with ApoTome.2 and a Camera Axiocam 702 mono. Specific filter sets were used for the visualization of green (Filter set 38 HE eGFP shift free [E] EX BP 470/40, BS FT 495, EM BP 525/50) and blue (Filter set 49 DAPI shift free [E] EX G 365, BS FT 395, EM BP 445/50) fluorescence. For genomic DNA extraction, fresh brain tissue was collected in RNAlater solution and kept at 4°C for 2 weeks. Then 250 μm thick sections containing the SCN were sliced using a microtome and the SCN region was microdissected under a fluorescent equipped stereomicroscope (Nikon SMZ-25).

### SCN slices and bioluminescence recording

Approximately 14 days later, following bilateral stereotactic injections, the mice were sacrificed and the SCN was dissected. Slices of 350 μm around the area of SCN were prepared with a tissue chopper between ZT4.8 and ZT6.3; 2 slices per animal were used. Slicing and recovery buffer contained of NMDG aCSF (85 mM NMDG, 9 mM MgSO4, 2.3mM KCl, 1.1 mM NaH_2_PO_4_, 0.5 mM CaCl_2_, 23 mM D-Glucose, 28 mM NaHCO_3_, 18 mM Hepes, 3 mM Na-pyruvate, 5 mM Na-ascorbate and 2 mM thiourea; pH 7.3-7.4; 300-310 mOsm/Kg according ^63^. Each slice was cultured in a single well of a 24-well plate in 300 μl of culture medium (0.7 x MEM Eagle medium with 1.7 mM MgSO_4_, 0.8 mM CaCl_2_, 11 mM D-Glucose, 17 mM NaHCO_3_, 25 mM Hepes, 0.4 mM GlutaMAX, 17% Horse serum, 0.8 mg/L Insulin, 0.8495 mM Ascorbic acid, 1% penicillin/streptomycin and 100 μM Luciferin; pH 7.3-7.4; 300-310 mOsm/Kg according ^63^). Viral infection and accurate injection localization of the SCN was evaluated with fluorescent imaging with THUNDER Imaging Systems widefield microscope (Leica) on the 8^th^ day in culture. Circadian bioluminescence was monitored by using photomultiplier tubes (PMTs) for approximately one week at 34.5 °C with 5% CO_2_ (in-house built device).

### Protein extraction and Western Blot

Total proteins from mouse liver were extracted in principle according to the NUN procedure ^64^. Freshly harvested liver extracts were homogenized in 2 tissue volume of Nuclear Lysis Buffer (10 mM Hepes pH 7.6, 100 mM KCl, 0.1 mM EDTA, 10% Glycerol, 0.15 mM spermine, 0.5 mM spermidine) for 20 seconds using a Teflon homogenizer. 4 tissue volumes of 2x NUN Buffer (2M Urea, 2% NP40, 0.6 M NaCl, 50 mM Hepes pH 7.6, 2 mM DTT, 0.1 mM PMSF and supplemented with complete protease inhibitor tablets, Roche) were added dropwise and on a vortex with constant low speed to ensure immediate mixing. The lysates were incubated on ice for 30 min and then cleared through centrifugation at 10000 rpm, 4°C, for 20 min. Supernatants were stored at -80°C. Aliquots of the lysates (20-30 µg of protein loaded per lane, either from a pool from 3 mice or from individual mice, as indicated) were separated by SDS-PAGE and transferred to PVDF membrane by dry transfer using an iBlot 2 gel transfer device. After blocking (5% milk in TBST; for 1 hour at room temperature), the membrane was incubated overnight at 4°C with appropriate dilutions of primary antibodies, including anti-CRY2 (kind gift from Ueli Schibler, Geneva), anti-ATF5 (Abcam-ab184923), and anti-HSP90 (Cell signaling-4874), p-eif2alpha (Cell signaling-9721), eif2alpha (Cell signaling-9722). Following TBST washing (3 × 5 minutes), the membranes were incubated with the appropriate secondary antibody conjugated with HRP for 60 minutes at room temperature, followed by washing as above. Chemiluminescence signal was detected with Supersignal West Femto Maximum Sensitivity Substrate (Thermo Fisher Scientific, 34095), as described by the manufacturer. The quantification of bands was performed using ImageJ software.

## Data and script availability

Data has been deposited at GEO (reviewer token). Computational scripts are accessible at:

## Acknowledgements

We thank Paul Franken and Yann Emmenegger for equipment and help with *in vivo* experiments, and Oliver Mühlemann for generous gift of SMG1i compound. Work in DG’s lab was funded by the University of Lausanne and by the Swiss National Science Foundation through the National Center of Competence in Research RNA & Disease (grant no. 141735) and through individual grant 179190; work in AK’s lab was funded by the Deutsche Forschungsgemeinschaft (DFG, German Research Foundation) - Project Number 278001972 - TRR 186.

## Supplementary material

**Supplementary Figure S1.**
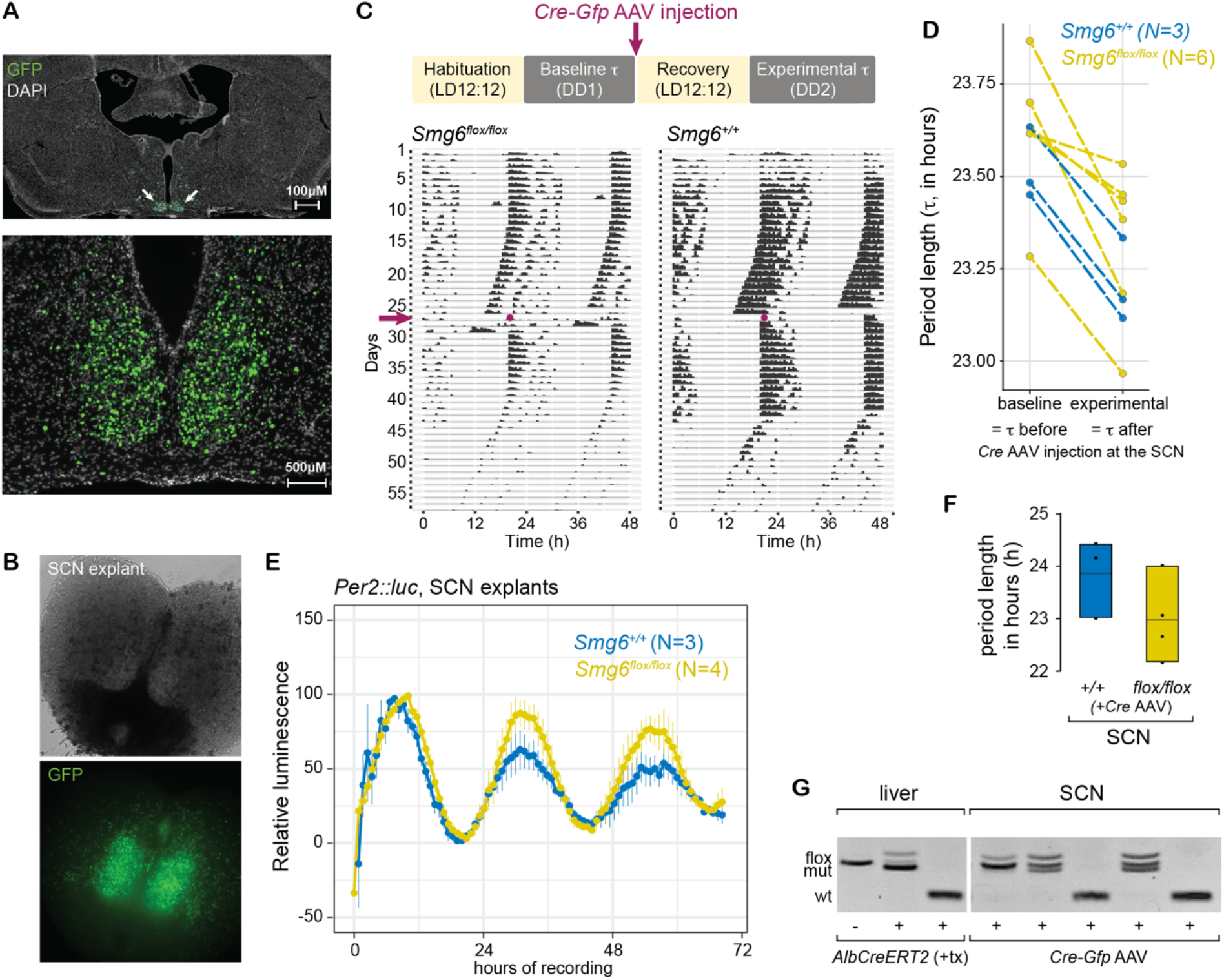
**A**. Representative microphotographs of SCN sections to assess effective targeting of the SCN. Viral expression can be estimated from GFP signal, encoded with Cre on the same virus. **B**. Same as A, but image taken during bioluminescence recording of SCN slices. **C**. Upper diagram: Rhythms of voluntary locomotor activity were recorded prior to and after the SCN injection of the Cre-and GFP-expressing AAV. Lower: Representative actograms of a *Smg6*^*flox/flox*^ and *Smg6*^*+/+*^mouse. The day of Cre::eGFP AAV injection is marekd by an arrow and a dot. **D**. Period lengths of circadian locomotor activity rhythms of *Smg6*^*flox/flox*^ (in yellow) and *Smg6*^*+/+*^ (in blue) mice before (DD1) and after (DD2) stereotaxic surgery. **E**. Averaged traces of *mPer::Luc* rhythms of AAV-injected *Smg6*^*flox/flox*^ (yellow) and *Smg6*^*+/+*^ (blue) SCN explants. **F**. Period lengths of *mPer::Luc* expression in AAV-injected *Smg6*^*flox/flox*^ (yellow) and *Smg6*^*+/+*^ (blue) SCN explants. **G**. Recombination efficiency following Cre induction was evaluated by genotyping of genomic DNA extracted from SCN slices (liver-specific mutants served as controls for the genotyping).

**Supplementary Figure S2:**
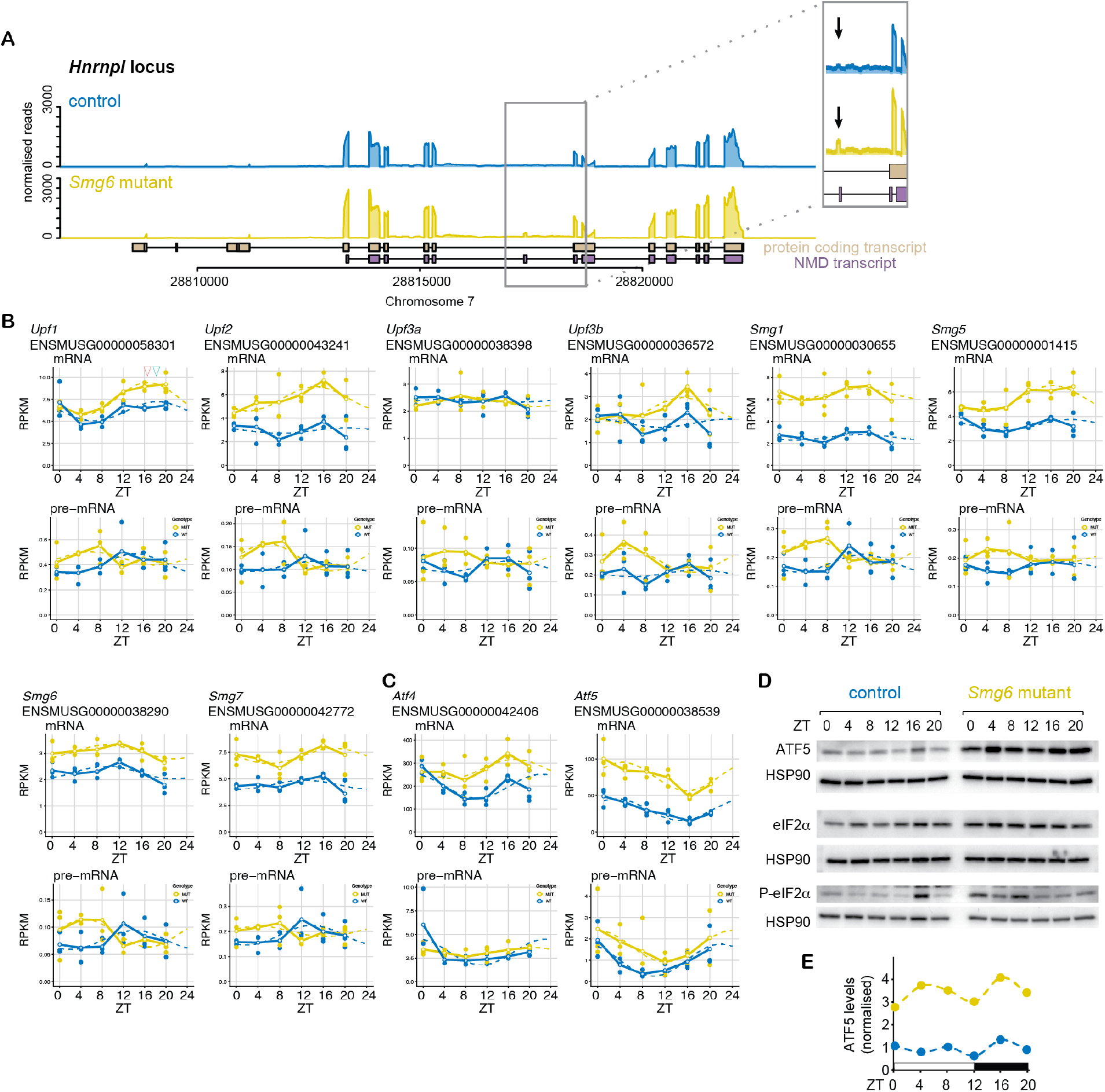
**A**. Read coverage on the *Hnrnpl1* locus indicates the specific upregulation of transcript isoforms that are NMD-annotated and that can be identified by specific exons (see arrows in insets) in liver tissue. **B**. RNA-seq data is plotted for indicated genes – that all encode components of the NMD machinery itself – for mRNA (upper panels; exonic reads) and pre-mRNA (lower panels; intronic reads) for *Smg6* mutants (yellow) and controls (blue). RPKM values of individual animals are shown as dots with solid lines connecting the means for each timepoint. The dashed lines represent the rhythmic data fit using the parameters from Metacycle. **C**. RNA-seq data is plotted for *Atf4* and *Atf5* for mRNA (upper panels; exonic reads) and pre-mRNA (lower panels; intronic reads) for *Smg6* mutants (yellow) and controls (blue). RPKM values of individual animals are shown as dots with solid lines connecting the means for each timepoint. The dashed lines represent the rhythmic data fit using the parameters from Metacycle. **D**. Western blot analysis of liver tissue (as in Fig. 4D) for ATF5, eIF2α and phospho-eIF2α in *Smg6* mutant and control liver samples; HSP90 served as loading control. **E**. Quantification of ATF5 signal, normalized to HSP90 as loading control, from Western blot shown in D.

**Supplementary Figure S3:**
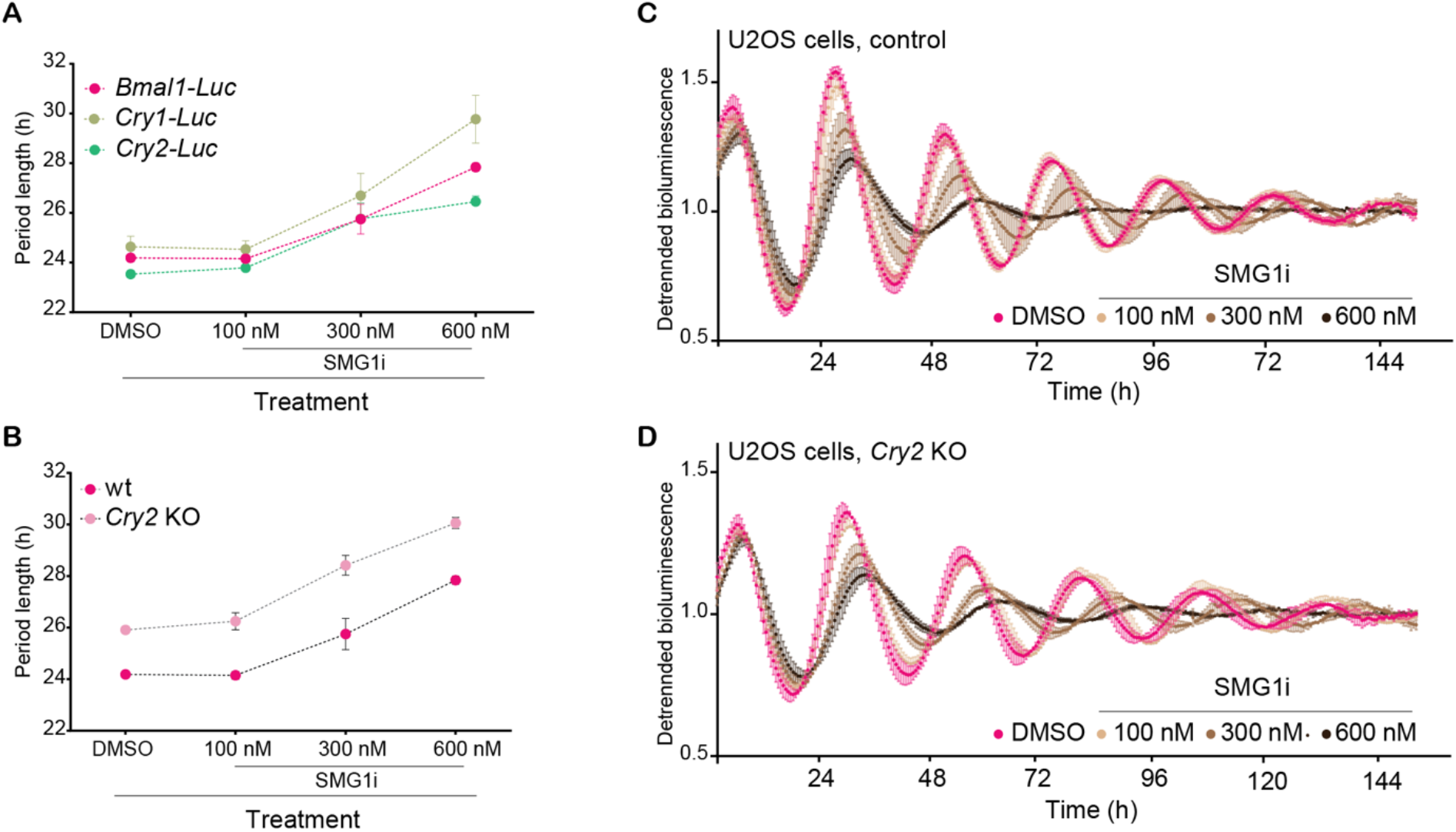
Pharmacological NMD inhibition prolongs circadian period in human osteosarcoma U2OS cells. **A**. Period length of the circadian reporters *Bmal1-Luc* (fuchsia), *Cry1-Luc* (khaki) or *Cry2-Luc* (green) in the presence of increasing concentrations of SMG1i or vehicle (DMSO, equal volume as for the highest SMG1i dose). **B**. Period length of the circadian reporter *Bmal1-Luc* in wt (pink) or *Cry2* KO (fuchsia) U2OS cells **C**. Traces of *Bmal1-Luc* detrended bioluminescence signal in wild-type U2OS cells treated with increasing dosage of SMG1i or vehicle. Solid circles represent mean, error bars represent standard deviation. **D**. Average traces of *Bmal1-Luc* detrended bioluminescence signal in *Cry2* KO U2OS cells treated with increasing concentrations of SMG1i or vehicle. Solid circles represent mean, error bars represent standard deviation.

## References

1. Karousis, E.D. & Muhlemann, O. Nonsense-Mediated mRNA Decay Begins Where Translation Ends. Cold Spring Harb Perspect Biol 11(2019).

2. Kurosaki, T., Popp, M.W. & Maquat, L.E. Quality and quantity control of gene expression by nonsense-mediated mRNA decay. Nat Rev Mol Cell Biol 20, 406–420 (2019).

3. Huth, M. et al. NMD is required for timely cell fate transitions by fine-tuning gene expression and regulating translation. Genes Dev 36, 348–367 (2022).

4. Boehm, V. et al. SMG5-SMG7 authorize nonsense-mediated mRNA decay by enabling SMG6 endonucleolytic activity. Nat Commun 12, 3965 (2021).

5. Mendell, J.T., Sharifi, N.A., Meyers, J.L., Martinez-Murillo, F. & Dietz, H.C. Nonsense surveillance regulates expression of diverse classes of mammalian transcripts and mutes genomic noise. Nat Genet 36, 1073–8 (2004).

6. Singh, G., Rebbapragada, I. & Lykke-Andersen, J. A competition between stimulators and antagonists of Upf complex recruitment governs human nonsense-mediated mRNA decay. PLoS Biol 6, e111 (2008).

7. Yepiskoposyan, H., Aeschimann, F., Nilsson, D., Okoniewski, M. & Muhlemann, O. Autoregulation of the nonsense-mediated mRNA decay pathway in human cells. RNA 17, 2108–18 (2011).

8. Karousis, E.D., Gypas, F., Zavolan, M. & Muhlemann, O. Nanopore sequencing reveals endogenous NMD-targeted isoforms in human cells. Genome Biol 22, 223 (2021).

9. Cox, K.H. & Takahashi, J.S. Circadian clock genes and the transcriptional architecture of the clock mechanism. J Mol Endocrinol 63, R93–R102 (2019).

10. Partch, C.L., Green, C.B. & Takahashi, J.S. Molecular architecture of the mammalian circadian clock. Trends in cell biology 24, 90–99 (2014).

11. Du, N.H., Arpat, A.B., De Matos, M. & Gatfield, D. MicroRNAs shape circadian hepatic gene expression on a transcriptome-wide scale. Elife 3, e02510 (2014).

12. Kojima, S., Sher-Chen, E.L. & Green, C.B. Circadian control of mRNA polyadenylation dynamics regulates rhythmic protein expression. Genes Dev 26, 2724–2736 (2012).

13. Kwon, Y.J., Park, M.J., Kim, S.G., Baldwin, I.T. & Park, C.M. Alternative splicing and nonsense-mediated decay of circadian clock genes under environmental stress conditions in Arabidopsis. BMC Plant Biol 14, 136 (2014).

14. Ri, H. et al. Drosophila CrebB is a Substrate of the Nonsense-Mediated mRNA Decay Pathway that Sustains Circadian Behaviors. Mol Cells 42, 301–312 (2019).

15. Wu, Y. et al. Up-Frameshift Protein UPF1 Regulates Neurospora crassa Circadian and Diurnal Growth Rhythms. Genetics 206, 1881–1893 (2017).

16. Glavan, F., Behm-Ansmant, I., Izaurralde, E. & Conti, E. Structures of the PIN domains of SMG6 and SMG5 reveal a nuclease within the mRNA surveillance complex. EMBO J 25, 5117–25 (2006).

17. Azzalin, C.M. & Lingner, J. The double life of UPF1 in RNA and DNA stability pathways. Cell Cycle 5, 1496–8 (2006).

18. Li, T. et al. Smg6/Est1 licenses embryonic stem cell differentiation via nonsense-mediated mRNA decay. EMBO J 34, 1630–47 (2015).

19. Gaidatzis, D., Burger, L., Florescu, M. & Stadler, M.B. Analysis of intronic and exonic reads in RNA-seq data characterizes transcriptional and post-transcriptional regulation. Nat Biotechnol 33, 722–9 (2015).

20. Koike, N. et al. Transcriptional architecture and chromatin landscape of the core circadian clock in mammals. Science 338, 349–54 (2012).

21. Janich, P., Arpat, A.B., Castelo-Szekely, V., Lopes, M. & Gatfield, D. Ribosome profiling reveals the rhythmic liver translatome and circadian clock regulation by upstream open reading frames. Genome Res 25, 1848–59 (2015).

22. French, C.E. et al. Transcriptome analysis of alternative splicing-coupled nonsense-mediated mRNA decay in human cells reveals broad regulatory potential. bioRxiv, 2020.07.01.183327 (2020).

23. Stratmann, M., Suter, D.M., Molina, N., Naef, F. & Schibler, U. Circadian Dbp transcription relies on highly dynamic BMAL1-CLOCK interaction with E boxes and requires the proteasome. Mol Cell 48, 277–87 (2012).

24. Brown, S.A., Zumbrunn, G., Fleury-Olela, F., Preitner, N. & Schibler, U. Rhythms of mammalian body temperature can sustain peripheral circadian clocks. Curr Biol 12, 1574–83 (2002).

25. Schuler, M., Dierich, A., Chambon, P. & Metzger, D. Efficient temporally controlled targeted somatic mutagenesis in hepatocytes of the mouse. Genesis 39, 167–72 (2004).

26. Yoo, S.-H. et al. PERIOD2:: LUCIFERASE real-time reporting of circadian dynamics reveals persistent circadian oscillations in mouse peripheral tissues. Proceedings of the National Academy of Sciences 101, 5339–5346 (2004).

27. Aschoff, J. & Pohl, H. Phase relations between a circadian rhythm and its zeitgeber within the range of entrainment. Naturwissenschaften 65, 80–4 (1978).

28. Granada, A.E., Bordyugov, G., Kramer, A. & Herzel, H. Human chronotypes from a theoretical perspective. PLoS One 8, e59464 (2013).

29. Katsioudi, G. et al. Recording of Diurnal Gene Expression in Peripheral Organs of Mice Using the RT-Biolumicorder. Methods Mol Biol 2482, 217–242 (2022).

30. Saini, C. et al. Real-time recording of circadian liver gene expression in freely moving mice reveals the phase-setting behavior of hepatocyte clocks. Genes & development 27, 1526–1536 (2013).

31. Hoekstra, M.M., Jan, M., Katsioudi, G., Emmenegger, Y. & Franken, P. The sleep-wake distribution contributes to the peripheral rhythms in PERIOD-2. Elife 10(2021).

32. Debruyne, J.P. et al. A clock shock: mouse CLOCK is not required for circadian oscillator function. Neuron 50, 465–77 (2006).

33. Kornmann, B., Schaad, O., Bujard, H., Takahashi, J.S. & Schibler, U. System-driven and oscillator-dependent circadian transcription in mice with a conditionally active liver clock. PLoS Biol 5, e34 (2007).

34. Hatano, M. et al. The 5’-untranslated region regulates ATF5 mRNA stability via nonsense-mediated mRNA decay in response to environmental stress. FEBS J 280, 4693–707 (2013).

35. Pakos-Zebrucka, K. et al. The integrated stress response. EMBO Rep 17, 1374–1395 (2016).

36. Wu, G., Anafi, R.C., Hughes, M.E., Kornacker, K. & Hogenesch, J.B. MetaCycle: an integrated R package to evaluate periodicity in large scale data. Bioinformatics 32, 3351–3353 (2016).

37. Castelo-Szekely, V. et al. Charting DENR-dependent translation reinitiation uncovers predictive uORF features and links to circadian timekeeping via Clock. Nucleic Acids Res 47, 5193–5209 (2019).

38. Gopalsamy, A. et al. Identification of pyrimidine derivatives as hSMG-1 inhibitors. Bioorg Med Chem Lett 22, 6636–41 (2012).

39. Langer, L.M., Bonneau, F., Gat, Y. & Conti, E. Cryo-EM reconstructions of inhibitor-bound SMG1 kinase reveal an autoinhibitory state dependent on SMG8. Elife 10(2021).

40. Zinshteyn, B., Sinha, N.K., Enam, S.U., Koleske, B. & Green, R. Translational repression of NMD targets by GIGYF2 and EIF4E2. PLoS Genet 17, e1009813 (2021).

41. Luck, S. & Westermark, P.O. Circadian mRNA expression: insights from modeling and transcriptomics. Cell Mol Life Sci 73, 497–521 (2016).

42. Hughes, M.E. et al. Guidelines for Genome-Scale Analysis of Biological Rhythms. J Biol Rhythms 32, 380–393 (2017).

43. Rey, G. et al. Genome-wide and phase-specific DNA-binding rhythms of BMAL1 control circadian output functions in mouse liver. PLoS Biol 9, e1000595 (2011).

44. Damiola, F. et al. Restricted feeding uncouples circadian oscillators in peripheral tissues from the central pacemaker in the suprachiasmatic nucleus. Genes Dev 14, 2950–61 (2000).

45. Miller, S. et al. Isoform-selective regulation of mammalian cryptochromes. Nature chemical biology 16, 676–685 (2020).

46. Anand, S.N. et al. Distinct and separable roles for endogenous CRY1 and CRY2 within the circadian molecular clockwork of the suprachiasmatic nucleus, as revealed by the Fbxl3Afh mutation. Journal of Neuroscience 33, 7145–7153 (2013).

47. Ukai-Tadenuma, M. et al. Delay in feedback repression by cryptochrome 1 is required for circadian clock function. Cell 144, 268–81 (2011).

48. Liu, A.C. et al. Intercellular coupling confers robustness against mutations in the SCN circadian clock network. Cell 129, 605–616 (2007).

49. Jaffrey, S.R. & Wilkinson, M.F. Nonsense-mediated RNA decay in the brain: emerging modulator of neural development and disease. Nature Reviews Neuroscience 19, 715–728 (2018).

50. Popp, M.W. & Maquat, L.E. Nonsense-mediated mRNA decay and cancer. Current opinion in genetics & development 48, 44–50 (2018).

51. Lindeboom, R.G., Supek, F. & Lehner, B. The rules and impact of nonsense-mediated mRNA decay in human cancers. Nature genetics 48, 1112–1118 (2016).

52. Ivanov, I., Lo, K., Hawthorn, L., Cowell, J.K. & Ionov, Y. Identifying candidate colon cancer tumor suppressor genes using inhibition of nonsense-mediated mRNA decay in colon cancer cells. Oncogene 26, 2873–2884 (2007).

53. Barmada, S.J. et al. Amelioration of toxicity in neuronal models of amyotrophic lateral sclerosis by hUPF1. Proceedings of the National Academy of Sciences 112, 7821–7826 (2015).

54. Kumar, M.S. et al. Dicer1 functions as a haploinsufficient tumor suppressor. Genes & development 23, 2700–2704 (2009).

55. Naviaux, R.K., Costanzi, E., Haas, M. & Verma, I.M. The pCL vector system: rapid production of helper-free, high-titer, recombinant retroviruses. Journal of virology 70, 5701–5705 (1996).

56. Stewart, S.A. et al. Lentivirus-delivered stable gene silencing by RNAi in primary cells. Rna 9, 493–501 (2003).

57. Salmon, P. & Trono, D. Production and titration of lentiviral vectors. Current protocols in human genetics 54, 12.10.1-12.10.24 (2007).

58. Dobin, A. et al. STAR: ultrafast universal RNA-seq aligner. Bioinformatics 29, 15–21 (2013).

59. Anders, S., Pyl, P.T. & Huber, W. HTSeq—a Python framework to work with high-throughput sequencing data. bioinformatics 31, 166–169 (2015).

60. Danecek, P. et al. Twelve years of SAMtools and BCFtools. Gigascience 10, giab008 (2021).

61. Love, M.I., Huber, W. & Anders, S. Moderated estimation of fold change and dispersion for RNA-seq data with DESeq2. Genome biology 15, 1–21 (2014).

62. Brancaccio, M. et al. Cell-autonomous clock of astrocytes drives circadian behavior in mammals. Science 363, 187–192 (2019).

63. Ting, J.T. et al. Preparation of acute brain slices using an optimized N-methyl-D-glucamine protective recovery method. JoVE (Journal of Visualized Experiments), e53825 (2018).

64. Lavery, D.J. & Schibler, U. Circadian transcription of the cholesterol 7 alpha hydroxylase gene may involve the liver-enriched bZIP protein DBP. Genes & development 7, 1871–1884 (1993).

